# Auxin and pectin remodeling interplay during rootlet emergence in white lupin

**DOI:** 10.1101/2021.07.19.452882

**Authors:** François Jobert, Alexandre Soriano, Laurent Brottier, Célia Casset, Fanchon Divol, Josip Safran, Valérie Lefebvre, Jérôme Pelloux, Stéphanie Robert, Benjamin Péret

**Affiliations:** BPMP, Univ Montpellier, CNRS, INRAE, Supagro, Montpellier, France; Umeå Plant Science Centre (UPSC), Department of Forest Genetics and Plant Physiology, Swedish University of Agricultural Sciences, Umeå, Sweden; UMR INRAE 1158 BioEcoAgro, BIOPI Biologie des Plantes et Innovation, SFR Condorcet FR CNRS 3417, Université de Picardie, Amiens, France

**Author notes:** Contributions: F.J. performed most of the experiments and analyzed results. A.S and F.J. analyzed RNAseq data, L.B. designed the binary vector pK7m24GW_CR for hairy root phenotyping experiments, C.C. and F.D. sampled and prepared RNA libraries for sequencing, J.S. generated *in silico* PG models, V.L. performed oligogalacturonide dosages, F.J., J.P. and V.L. analyzed OG dosages, F.J., S.R and B.P. designed the research, F.J, S.R and B.P. wrote the article.

## Abstract

Secondary root emergence is a crucial trait that shapes the plant’s underground system. Virtually every developmental step of root primordium morphogenesis is controlled by auxin. However, how the hormone controls cell separation in primordium-overlaying tissues through wall loosening is poorly understood. Here, we took advantage of white lupin and its spectacular cluster root development to assess the contribution of auxin to this process. We show that auxin’s positive role on rootlet emergence is associated with an upregulation of cell wall pectin modifying and degrading genes. Downregulation of a pectinolytic enzyme gene expressed in cells surrounding the primordium resulted in delayed emergence. Pectins were demethylesterified in the emergence zone and auxin treatment further enhanced this effect. Additionally, we report specific rhamnogalacturonan-I modifications during cortical cell separation. In conclusion, we propose a model in which auxin has a dual role during rootlet emergence: Firstly, through active pectin demethylesterification and secondly by regulating the expression of cell wall remodeling enzymes.

## Introduction

Climate change and extensive use of modern agricultural systems has brought to light alarming issues for crop production. In addition to weather hazards, farmers must deal with impoverished soils while maintaining a sustainable yield. Thus, plants need to be carefully selected based on their ability to cope with specific stresses. White lupin (*Lupinus albus*) is a nitrogen-fixing annual plant from the Leguminosae family that remarkably thrives on phosphorus poor soils. As an adaptation to low phosphate conditions, white lupin grows specialized root structures called cluster roots. These secondary roots almost synchronously develop hundreds of rootlets, defined by their short life and determinate growth in a brush-like structure (Vance *et al.*, 2003). Rootlet morphogenesis in lupin differs from the simplified, however peculiar, lateral root development of *Arabidopsis thaliana*. In contrast to *Arabidopsis*, where the lateral root primordium derives exclusively from mitotically activated pericycle cells, several tissue layers can contribute to the lupin rootlet primordium (Gallardo *et al.*, 2018), which is in common with many angiosperms (Xiao *et al.*, 2019). Rootlet primordia cross these barriers successfully without damaging the outer tissues despite their large number and proximity.

During the growth of the rootlet primordium, the surrounding cells are subjected to morphological adjustments including cell division and detachment. Yet, the plant cell wall is an obstacle to the latter, acting as a glue between cells. The cell wall is composed of polysaccharides including cellulose, hemicellulose and pectins together with structural proteins. This dynamic “frame-like” structure provides support and protection but is flexible enough to accommodate the cell fate. The phytohormone auxin is one of the factors regulating wall loosening, notably through the modification of gene expression (Majda et Robert, 2018). During *Arabidopsis* lateral root emergence, an auxin maximum is generated by the sequential activation of the auxin efflux carrier PIN-FORMED3 (PIN3) and influx carrier AUXIN TRANSPORTER-LIKE PROTEIN 3 (LAX3) in the primordium-overlaying cells (Swarup *et al.*, 2008, Péret *et al.*, 2013). This auxin sink formation induces the expression of the cell wall remodeling enzymes *XTH23/XYLOGLUCAN ENDOTRANSGLYCOSYLASE6* (*XTR6*) and *POLYGALACTURONASE INVOLVED IN LATERAL ROOT* (*PGLR*) (Swarup *et al.*, 2008), two carbohydrate-active enzymes affecting wall integrity and reducing cell adhesion. The INFLORESCENCE DEFICIENT IN ABSCISSION (IDA) peptide and its receptor-like kinases HAESA/HAESA-LIKE2 (HAE/HSL2) belong to this auxin-dependent network and regulate the expression of *XTR6* and *PGLR* through a recently identified MITOGEN-ACTIVATED PROTEIN KINASE cascade (Kumpf *et al.*, 2013, Zhu *et al.*, 2019). Cell wall remodeling enzyme activity can be modulated by REACTIVE OXYGEN SPECIES (ROS)-mediated cell wall acidification, as a consequence of the spatial activation of *RESPIRATORY BURST OXIDASE HOMOLOG* (*RBOH*) genes by auxin (Orman-Lizega *et al.*, 2016). In addition, mutants with altered cell wall composition show modified root architecture (Roycewicz and Malamy, 2014) and the mechanical properties of overlaying tissues influence lateral root primordium development (Lucas *et al.*, 2013). A recent report revealed the importance of pectin homogalacturonan methylesterification to regulate pre-branching site formation along the *Arabidopsis* primary root (Wachsman *et al.*, 2020). The authors also suggest that methylesterification in primordium-overlaying cell walls could play a role in facilitating their emergence. However, the modifications of cell wall chemistry and their consequences during lateral root emergence are still unclear.

Taking advantage of white lupin cluster roots and their perfectly synchronized rootlet emergence, we assessed the contribution of auxin to cell wall modifications during this process. In this study, we found a positive effect of auxin on rootlet emergence with transcriptomic signatures associated with oxidative stress, transcriptional regulation and cell wall remodeling. Downregulation of an auxin-responsive polygalacturonase gene expressed in the cell layer surrounding the rootlet primordium delayed emergence, suggesting the relevance of localized pectin degradation. Interestingly, homogalacturonans, the known target of polygalacturonases, were strongly demethylesterified in the emergence zone with auxin enhancing this modification. Yet, the methylesterification degree of homogalacturonan did not vary in cortical cells whether challenged by an emerging rootlet primordium or not. Intriguingly, rhamnogalacturonan-I (1,4)-β-D-galactans and extensin glycoproteins showed differential distribution and could be a hallmark of cell separation and/or mechanically challenged cells. In summary, we suggest a model in which auxin acts before rootlet primordium emergence by reducing methylesterification of pectins and thus priming the cell wall for the sequential action of pectin degradation enzymes.

## Results

### Rootlet primordium emergence is positively regulated by auxin

Auxin is a central regulator in lateral root primordium initiation and emergence in higher plants (Du et Scheres, 2018). In white lupin, the natural auxin indole-3-acetic acid (IAA) has been shown to induce the formation of cluster roots under phosphate deficiency (Neumann *et al.*, 2000, Meng *et al.*, 2012). To understand the precise contribution of auxin in the white lupin rootlet emergence process, we tested the effect of IAA on cluster root development. IAA exogenous application modified the root system architecture quite drastically (Fig. 1a, 1b, Extended Data Fig. 1). Primary and secondary root elongation, including that of specialized cluster roots, was reduced in a dose dependent manner from 1 μM IAA with a concomitant decrease in rootlet primordium number, while the overall rootlet density was unchanged (Fig. 1b). We assigned the pre-emerged rootlet primordia to seven developmental stages (Gallardo *et al.*, 2018). The analysis of rootlet primordium stage distribution revealed that 70% were emerged at the highest concentration of IAA compared to only 30% in the control conditions (Fig. 1c). Accordingly, less primordia were found in stages I to VI in IAA treated plants compared to control plants. Overall, these data suggest that IAA modifies the lupin root architecture by promoting primordium emergence in a dose dependent fashion. To further understand how auxin influences lupin root architecture, we also challenged the plants with naphthylphthalamic acid (NPA) (Abas *et al.*, 2021) and 1-naphthoxyacetic acid (1-NOA) (Lankova *et al.*, 2010), auxin efflux and influx inhibitors respectively (Fig. 1a, Extended Data Fig. 1). NPA treatment strongly decreased rootlet number and density (Fig. 1b) and delayed rootlet primordium emergence (Fig. 1c). Inhibition of auxin influx machinery with 1-NOA had a mild negative effect on rootlet number but did not affect rootlet density (Fig. 1b). However, at 25 μM 1-NOA, emergence was delayed as shown by a higher number of primordia in the intermediate stages V-VI (Fig. 1c). Overall, these data demonstrate an essential role of polar auxin transport in rootlet primordium morphogenesis while auxin uptake has a more specific role during primordium emergence. Auxin modulates the expression of a plethora of downstream target genes. To determine the extent of auxin transcriptional responses during rootlet emergence, the *DR5::nlsYFP* synthetic auxin reporter was expressed transiently in lupin hairy roots. A strong activation of *DR5::nlsYFP* expression was found in the pericycle cell layer of cluster roots (Fig. 1d). We also detected a weak *DR5::nlsYFP* signal at the tip of the rootlet primordium starting from stage V. These results suggest that auxin transcriptional responses in secondary root primordia are mostly conserved between lupin and *Arabidopsis*.

**Figure 1:**
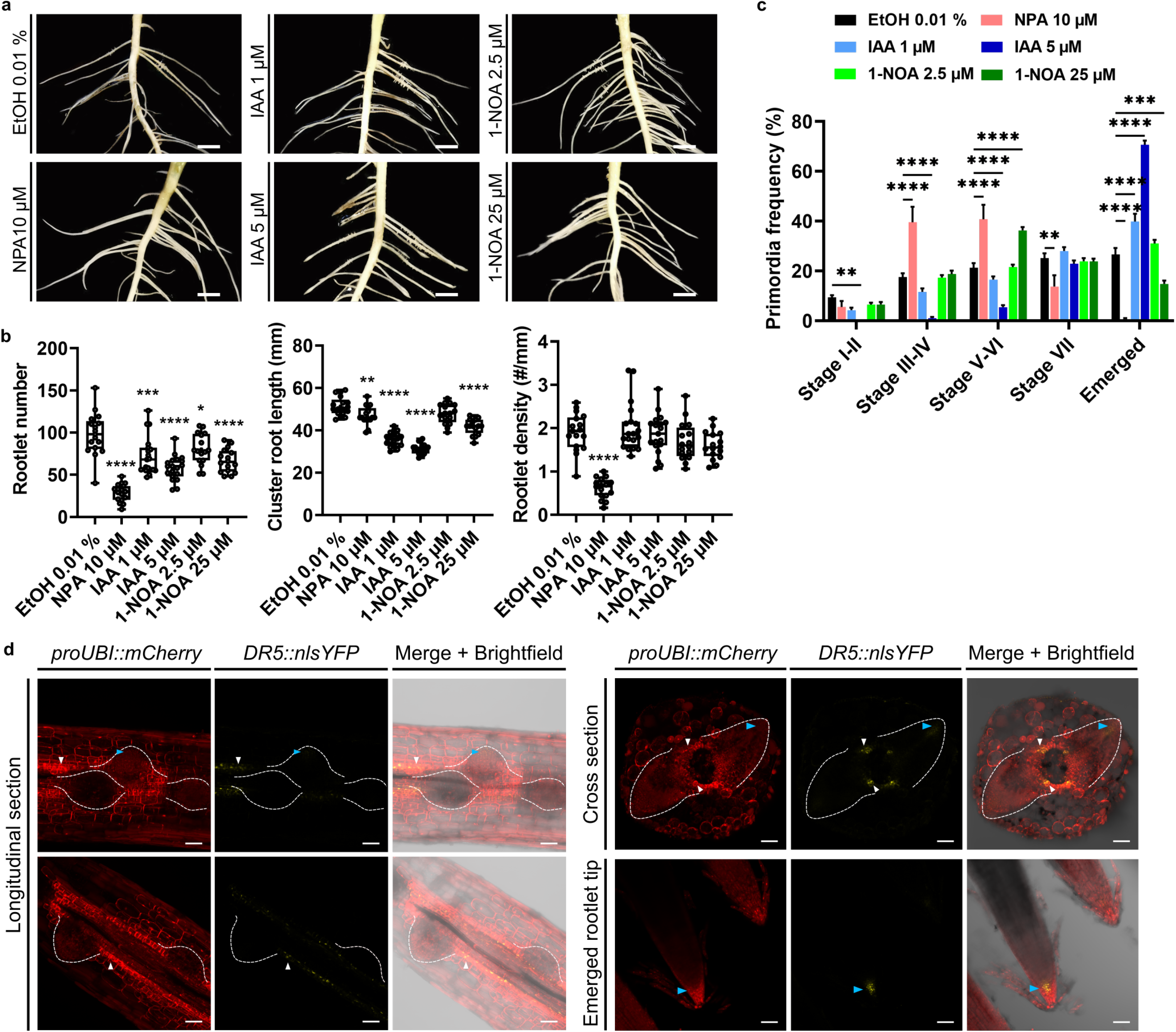
Auxin regulates rootlet development and primordium emergence. **a**, Representative pictures of the cluster roots from nine day-old *L. albus* grown in hydroponics medium after two days supplemented with 0.01% ethanol (control treatment), 10 μM NPA, 1 μM IAA, 5 μM IAA, μM 1-NOA or 25 μM 1-NOA. Bar scale: 1 cm. **b**, Rootlet number, cluster root length and rootlet density in the 4 upper cluster roots in plants treated with 0.01% ethanol (n = 17), 10 μM NPA (n = 16), 1 μM IAA (n = 19), 5 μM IAA (n = 19), 2.5 μM 1-NOA (n = 16) or 25 μM 1-NOA (n = 16). Statistical significance (compared to control) was computed by the Dunnett multiple comparison test: ****: pVal < 0.001, ***: pVal < 0.005, **: pVal < 0.01. **c**, Frequency of primordium stages found in the 4 upper cluster roots in plants treated with 0.01% ethanol (black bars, n = 17), 10 μM NPA (pink bars, n = 16), 1 μM IAA (light blue bars, n = 19), 5 μM IAA (dark blue bars, n = 19), 2.5 μM 1-NOA (light green bars, n = 16) or 25 μM 1-NOA (dark green bars, n = 16). Statistical significance (compared to control) was computed by the Dunnett multiple comparison test: ****: pVal < 0.001, ***: pVal < 0.005, **: pVal < 0.01, *: pVal < 0.05. **d**, Synthetic auxin response reporter DR5 is active in the pericycle cell layer of the cluster root emergence zone and at the tip of rootlet primordia. Free-hand sections were done immediately after harvesting fresh transformed roots. White dashed lines outline rootlet primordia, white and blue arrows respectively indicate pericycle cell layer and the tip of a primordium/rootlet. Scale bars for longitudinal sections: 50 μm. Scale bars for cross sections and emerged rootlet tips: 100 μm.

### Auxin-treated cluster root transcriptome identifies intense cell wall remodeling occurring during rootlet emergence

To better understand the effect of auxin on rootlet primordium emergence (Fig. 1c), we performed RNA sequencing on cluster root segments corresponding to the rootlet emergence zone (Fig. 2a). We collected samples treated with IAA at early (30 min, 1 and 2 hours) and late time points (6, 12, 24 and 48 hours) after treatment to assess the corresponding auxin-induced transcriptomic regulation. In total, 856 differentially expressed genes (DEG) compared to control treatment were detected and classified into seven hierarchical clusters based on their expression patterns (Fig. 2b, Supplementary Table 1). Three expression clusters gathered most of the DEG with 248 (cluster 1), 317 (cluster 2) and 141 (cluster 3) DEG and could be defined by distinct transcriptomic signatures (Fig. 2c). The remaining DEG were grouped into four minor clusters showing oscillating responses (Extended Data Fig. 2). We then conducted a gene ontology enrichment analysis to enable the functional interpretation of those clusters (Fig. 2c, Extended data Fig. 2, Supplemental Table 2). Cluster 1 included genes that were slightly upregulated by auxin in the early phase and then downregulated at the late time points. Genes in this cluster were mainly associated with oxidative stress-related processes with the identification of several putative peroxidase encoded genes. Interestingly, ROS production and signaling have been recently proposed as positive regulators of lateral root emergence in *Arabidopsis* (Manzano *et al.*, 2014, Orman-Lizega *et al.*, 2016). Cluster 2 was assigned to early auxin upregulated genes for which the expression peaked after 30 minutes of treatment and returned gradually to a low level after that (Fig. 2c). Those early-auxin induced genes were associated to processes such as defense responses and DNA-binding transcription factor activity. Cluster 3 differed by its expression and was defined by genes showing a late and steady auxin-induced upregulation starting after two hours of treatment (Fig. 2c). Remarkably, this cluster was enriched in genes with predicted roles in carbohydrate metabolic process, lyase and polygalacturonase activities. This suggests a prominent action of auxin on quick but transient transcriptional responses, rapid modulation of oxidative stress and most importantly a slow but durable cell wall remodeling mechanism occurring during rootlet emergence.

**Figure 2:**
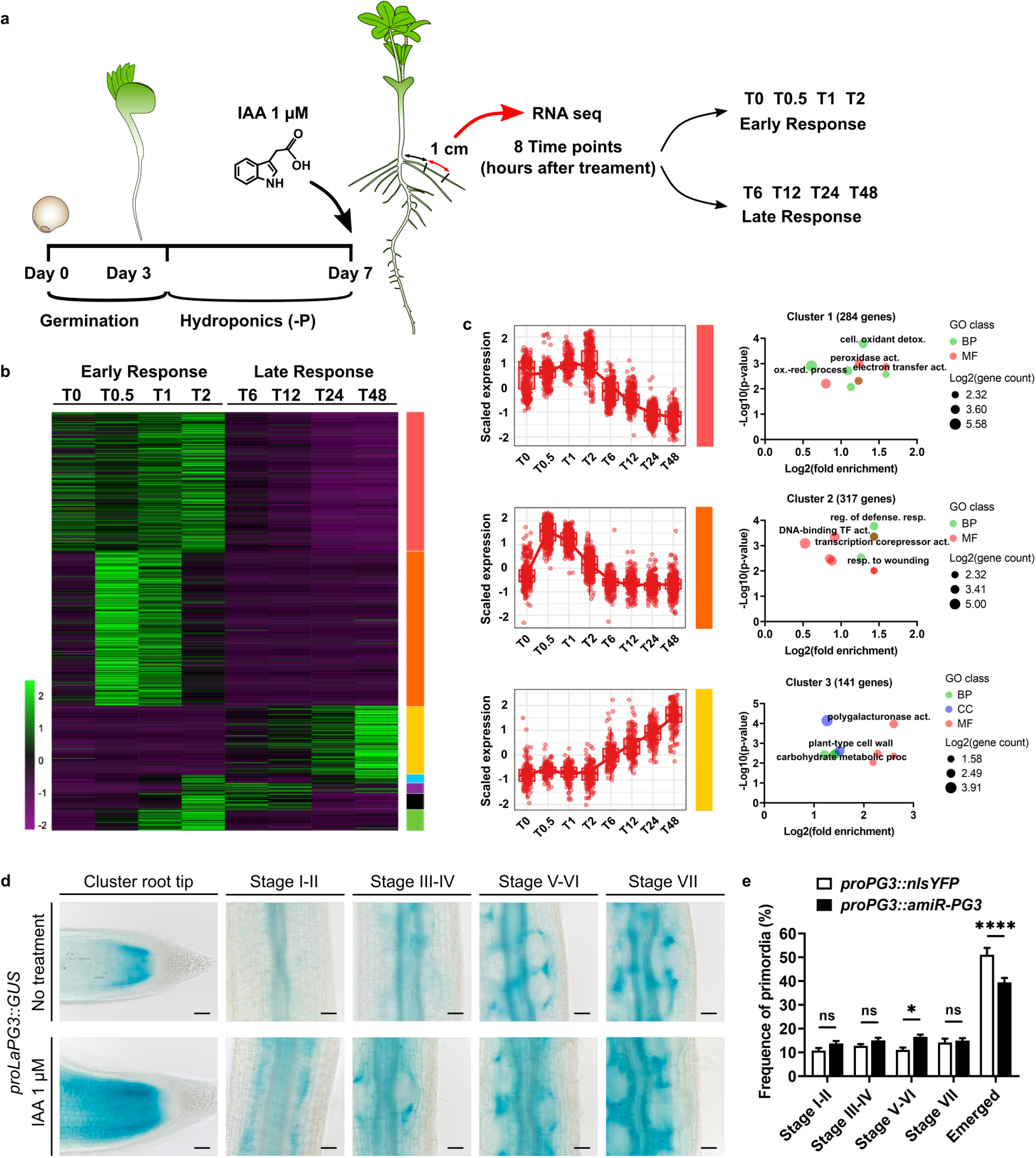
Auxin transcriptome landscape identifies cell wall related genes linked to rootlet emergence. **a**, Auxin RNA sequencing experiment overview. Lupin seeds were germinated in vermiculite for three days before the seedlings (“mohawk” stage) were transferred to hydroponic medium without phosphate (-P) to induce the formation of cluster roots (Day 3). Four days later (Day 7), 1 μM IAA was added to the hydroponic medium and 1 cm segments of cluster roots 1 cm distant from the primary root, corresponding to the emergence zone, were harvested for RNA extraction and further RNA sequencing at 8 different time points: T0 (just before the treatment), half an hour (T0.5), one hour (T1) and two hours (T2) after the treatment to assess the early auxin transcriptomic responses and six hours (T6), 12 hours (T12), one day (T24) and two days (T48) later for the late auxin transcriptomic responses. **b**, Heatmap of white lupin auxin-regulated transcripts. Normalized expression levels are shown as a z-score (See Material and Methods section for further details) and hierarchical clusters of the differentially expressed genes (DEG) are displayed by the colored squares on the right of the heatmap. Expression patterns for each cluster are shown in Extended data Figure 2. **c**, the three largest expression clusters (red, orange and yellow) show distinct gene expression patterns and are enriched in specific gene ontology terms. GO terms of significant importance are shown (red circle: molecular function, MF, green circle: biological process, BP, blue circle: cellular component, CC). **d**, proLaPG3::GUS localization in cluster root tips and at different rootlet primordium development stages in regular hydroponic medium (top) and after 48 hours of 1 μM IAA treatment (bottom). Scale bars: 100 μm. **e**, frequency of primordium stages in cluster roots from hairy root composite plants expressing proPG3::amiR-PG3 (n = 27 roots) and proPG3::nlsYFP (n = 28) as a control. Statistical significance was computed by the Šídák’s multiple comparisons test. *: pVal < 0.05, ****: pVal < 0.001, ns: non-significant.

### Pectin remodeling genes show specific expression pattern during rootlet emergence

Cell wall properties of primordium-overlaying tissues greatly impact the emergence process. Previous studies discovered a large subset of cell wall remodeling genes involved in lateral root emergence in *Arabidopsis*. We established a list of orthologs for important known regulators of cell wall loosening during lateral root morphogenesis including the closest lupin orthologs of *EXPANSIN A1* (EXPA1 - *At1g69530*, Ramakrishna *et al.*, 2019), *EXPANSIN A17* (*EXPA17* - *At4g01630*, Lee et Kim, 2013), *XYLOGLUCAN ENDOTRANSGLYCOSYLASE 6* (*XTR6*/*XTH23* - *At4g25810*, Swarup *et al.*, 2008), homogalacturonan degrading enzymes *POLYGALACTURONASE LATERAL ROOT* (*PGLR* - *At5g14650*, Kumpf *et al.*, 2013) and *PECTIN LYASE A2* (*PLA2 - At1g67750*, Swarup *et al.*, 2008) and two homogalacturonan modifying enzymes *PECTIN METHYLESTERASE 3* and *PECTIN METHYLESTERASE INHIBITOR 9* (*PME3* - *At3g14310* and *PMEI9* - *At1g62770*, Hocq *et al.*, 2017a) (Extended Data Fig. 3a). Next, we assessed the expression patterns of the selected candidates in lupin hairy roots transformed with promoter::GUS fusion constructs (Extended Data Fig. 3b and Extended Data Fig. 4). Some of the selected candidate genes that were not identified in our transcriptomic dataset displayed overlapping gene expression patterns. *LaXTR1* and *LaPME1* were expressed in the cluster root tip and rootlet primordium, while expression of *LaEXP1* and *LaPMEI1* were restricted to the primordium base and pericycle cells (Extended Data Fig. 3b). Interestingly, *proLaPMEI2::GUS* signal was found in the pericycle, endodermis and inner cortex surrounding rootlet primordia. In order to better understand the link between auxin and cell wall modification during rootlet emergence, we analyzed the expression pattern of additional auxin responsive genes present in cluster 3 (Extended Data Fig. 4). *O-GLYCOSYL HYDROLASE FAMILY 17L* (*LaOGH17L*) and *TRICHOME BIREFRINGENCE-LIKE 1* (*LaTBRL1*) were expressed in the cluster root at the base of the rootlet primordium and in the elongation zone (visible in stages VII-VIII). *PECTIN LYASE-LIKE 1* (*LaPLL1*) was expressed specifically in young vascular tissues (stages V – onward). *ProLaPLL2::GUS* displayed a broad signal in the meristematic zone of the cluster root and rootlet primordium. The galactose binding protein *DOMAIN OF UNKNOWN FUNCTION 642* gene (*LaDUF642*) was expressed in the elongation zone but also in the newly formed epidermis at the tip of the cluster root and rootlet primordium (stages V – onward) while *LaEXP2* expression was restricted to the epidermis in the elongation zone. *ProLaPG1::GUS* and *proLaPG2::GUS* activity were found in the vasculature including pericycle cells and *proLaPG2::GUS* was additionally found in the elongation zone of the emerging rootlet primordium (stages VII-VIII).

### Auxin responsive *Lupinus albus POLYGALACTURONASE3* (*LaPG3*) is expressed in outer cortex cells overlaying rootlet primordium

Despite being expressed at a low basal level, a third *PG* gene was highly responsive to auxin treatment (*LaPG3* – Extended Data Fig. 5). The LaPG3 protein sequence presents a high percentage of identity (60%) when compared to its closest ortholog in *Arabidopsis*, POLYGALACTURONASE INVOLVED IN LATERAL ROOT (AtPGLR) (Extended Data Fig. 6a). Structural modeling of LaPG3 showed that it contains a common right-handed β-parallel fold characteristic for pectinases, including fungal endo-PGs and exo-PGs (Extended Data Fig. 6b). Structural alignment with published PG structures (*Pectobacterium carotovorum*, *Aspergillus niger* and *Aspergillus aculeatus*) confirmed that LaPG3 is likely to act as an endo-PG, with conserved amino acid motifs at the active site including Asn211-Thr212-Asp213 (NTD), Asp234-Asp235 (DD), Gly256-His257-Gly258 (GHG) and Arg291-Ile292-Lys293 (RIK) (Extended Data Fig. 6c and d). The activity of its promoter was clearly visible in cluster root tips and even more striking in primordium-overlaying cells, excluding the endodermis, starting from stages III-IV (Fig. 2d). *ProLaPG3::nlsYFP* transformed roots confirmed the expression of the fluorescent protein in cells that undergo separation (Extended Data Fig. 7a). Interestingly, no signal was found in the mitotically activated endodermis and inner cortex cells. IAA treatment confirmed the responsiveness of *LaPG3* to the hormone, expanding the signal of *proLaPG3::GUS* to the external cortex (Fig. 2d). These results suggest that spatial regulation of auxin responsive *LaPG3* expression is important for rootlet primordium emergence. On top of that, GUS coloration was not found in the epidermis suggesting a different mechanism affecting cell separation in this layer. To understand the functional role of *LaPG3*, we expressed an artificial micro-RNA targeting its transcript under the control of the native promoter (*proLaPG3::amiR-LaPG3*) in lupin hairy roots (Extended Data Fig. 7b). While cluster root length, rootlet number and density were not affected in hairy roots expressing *proLaPG3::amiR-LaPG3* compared to the control *proLaPG3::nlsYFP* (Extended Data Fig. 7c), rootlet primordium emergence was slightly delayed in *proLaPG3::amiR-LaPG3* cluster roots (Fig. 2e). Gene expression analysis showed a 4-fold downregulation of *LaPG3* in *proLaPG3::amiR-LaPG3* cluster roots (Extended Data Fig. 7d). Furthermore, the expression of the auxin-responsive *GRETCHEN-HAGEN3* (*GH3*) genes *LaGH3.3*, *LaGH3.5* and *LaGH3.6* were increased in *proLaPG3::amiR-LaPG3* cluster roots, possibly reflecting a feedback between *LaPG3* and auxin homeostasis gene expression (Extended Data Fig. 7d).

### Oligogalacturonide profiling reveals auxin-induced pectin remodeling in cluster roots

Homogalacturonans (HG) are secreted into the cell wall as a polymer of galacturonic acids (GalA) which can be methylesterified and/or acetylated. It is suggested that the degree of methylesterification (DM) or acetylation (DA) of the HG plays a role in the availability of substrate for degradation enzymes such as polygalacturonases (Hocq *et al.*, 2017b). To understand the contribution of auxin-mediated changes in HG structure during rootlet emergence, we used an LC-MS/MS oligogalacturonide profiling method to reveal changes in HG composition in our cluster root samples (Voxeur *et al.*, 2019, Hocq *et al.*, 2020). For that purpose, we harvested segments corresponding to the “emergence zone” (as previously done for RNAseq) and the distal part of the cluster root, referred as the “CR tip zone” (Fig. 3a). Using *Aspergillus* PG, we then performed oligogalacturonide fingerprinting on these segments at T0, T24 hours and T48 hours after auxin treatment. Most of the oligogalacturonides released from the tip zone displayed a low degree of polymerization (DP ≤ 3) (Fig. 3b), which slightly accumulated over time in correlation with a decrease in high DP oligogalacturonides (DP ≥ 4) (Fig. 3b). Principal component analysis revealed a strong effect of auxin treatment in the accumulation of low DP oligogalacturonides (GalA2 and GalA3) compared to mock conditions. Accordingly, oligogalacturonides of high DP (DP ≥ 4), although weakly represented, were reduced following auxin treatment (Fig. 3b). Those effects were visible after 24 hours of treatment and stable after 48 hours (Fig. 3b). In contrast, oligogalacturonides that were released from the emergence zone differed over the time course. For example, although GalA2 was weakly represented at T0 (< 1% of total oligogalacturonides) its proportion was significantly increased (reaching 10%) at T24 (Fig. 3c). In the meantime, high DP oligogalacturonides (DP ≥ 3) were less abundant in the total released fraction at T24 and T48. However, auxin treatment had only a moderate impact on the DP of total oligogalacturonides released in the emergence zone and we observed a slight increase of GalA3 fraction while the GalA4 fraction was diminished (Fig. 3c). We next examined the DM and DA of the oligogalacturonides from the two segments of interest (Fig. 3d, e). In the cluster root tip zone, only slight changes in methylesterification or acetylation degrees occurred during the time course in control conditions, but auxin induced a strong demethylesterification of GalA species at T24 (Fig. 3d). A transient auxin-induced increase in acetylated oligogalacturonides compared to the control was observed at T24 before stabilizing to the control levels at T48 (Fig. 3d). This suggests a robust and early effect of auxin on HG modifications in the first 24 hours following the treatment in the CR tip zone. In the emergence zone, the already low level of released methylesterified oligogalacturonides was stable over time but was further reduced following auxin treatment after 24 hours (Fig. 3e). Remarkably, the proportion of acetylated oligogalacturonides diminished during the control time course but was not affected by auxin (Fig. 3e) suggesting an important relation between early rootlet emergence events and acetylation of HG in control conditions. We briefly summarized our findings in the model shown in Fig. 3f. When the plants are grown in control conditions, the tip zone of the cluster root is not subjected to dramatic changes in term of pectin modifications such as HG methylation. A brief variation of acetylation status was observed after 24 hours but stayed stable over time. The DP of oligogalacturonides released was fairly high. However, in the emergence zone in control conditions, we observed a low DM from the beginning of the time-course (T0) while the DA decreased substantially after 24 hours. Likewise, oligogalacturonide DP, which was lower than in the tip zone, decreased after 24 hours. When IAA treatment was applied at T0, this affected oligogalacturonide DM, which was dramatically lower after auxin treatment than in the control conditions for both zones. Meanwhile, upon auxin treatment, oligogalacturonide DA remained stable in both zones while DP was reduced in the tip zone similarly to the untreated emergence zone.

**Figure 3:**
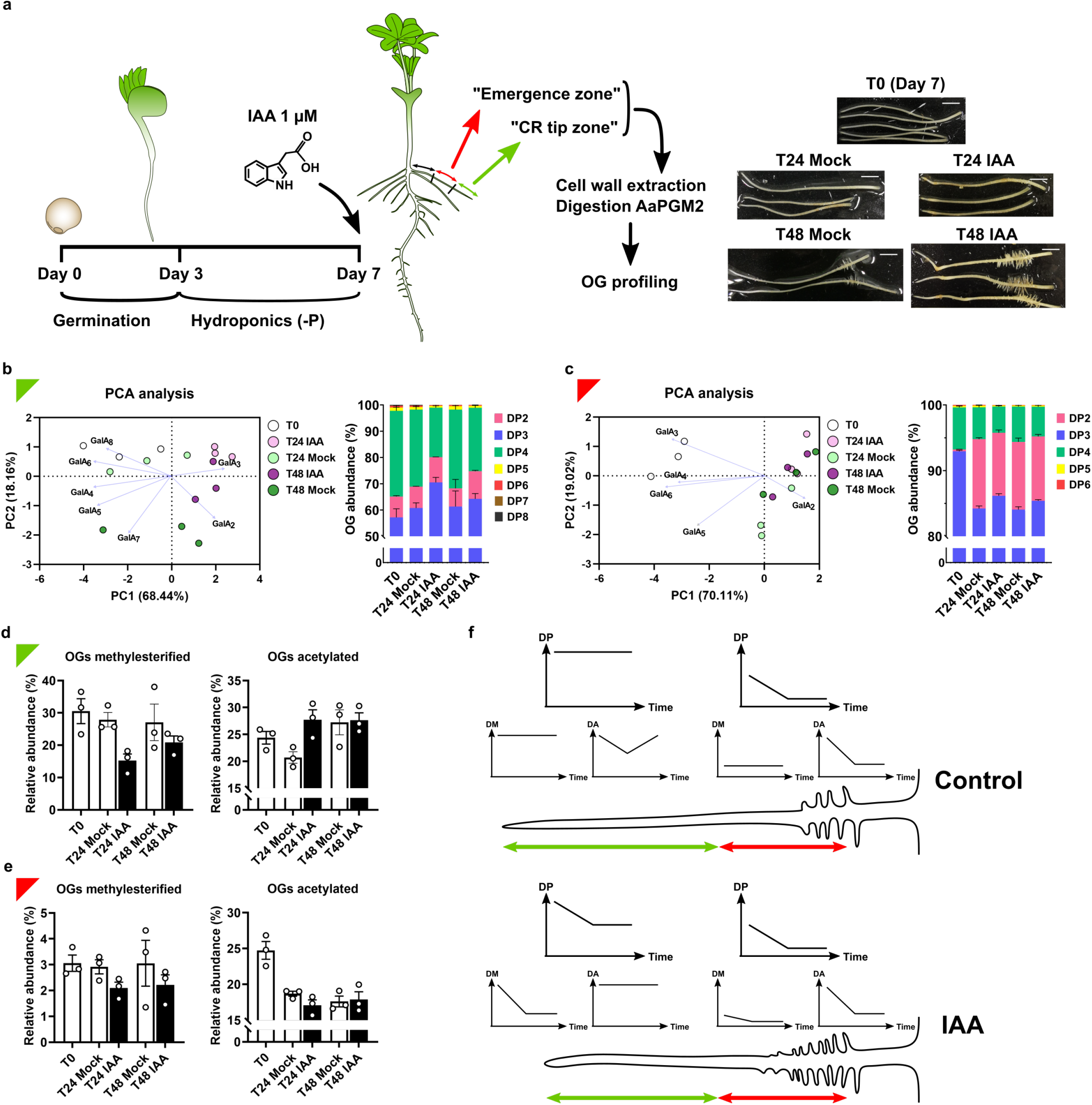
Oligogalacturonide profiling reveals auxin-induced pectin remodeling in cluster roots. **a**, Auxin oligogalacturonide (OG) profiling experiment overview. Lupin seeds germinated in vermiculite for three days before the seedlings (“mohawk” stage) were transferred to hydroponic medium without phosphate (-P) to induce the formation of cluster roots (Day 3). Four days later (Day 7), 1 μM IAA or 0.01% ethanol (Mock) were added to the hydroponic medium and 1 cm segments of cluster roots 1 cm distant from the primary root (hereafter named emergence zone, red color code) and the distal part of the cluster root (CR tip zone, green color code) were harvested for cell wall extraction, digestion with *Aspergillus aculeatus* endo-polygalacturonase M2 (AaPGM2) and OG profiling according to Voxeur *et al.*, 2019. The samples shown in the pictures (bars: 0.5mm) were dissected and collected at T0 (just before the auxin treatments), and at one day (T24) and two days (T48) after auxin treatments. **b**, Effect of the auxin treatment on GalAx species released in the "CR tip zone" during the time course previously described. Left plot: bi-dimensional plot of principal components calculated by performing PCA of the different OG species relative abundances (grouped by degree of polymerization) for each treatment and time point (3 replicates each). Vectors describe the contribution of the OG species to the biplot. Right plot: OG abundance (% of total OG detected) in the CR tip zone at T0 and 24 and 48 hours after treatment with ethanol (T24 and T48 Mock) or IAA (T24 and T48 IAA). **c**, Effect of the auxin treatment on GalAx species released in the "emergence zone" during the time course previously described. See **b** for descriptions of the plots. **d**, Relative abundance of methylesterified OG (GalAxMemAcn, x, m ≥ 1, n ≥ 0) or acetylated OG (GalAxMemAcn, x, m ≥ 0, n ≥ 1) from the CR tip zone. **e**, Relative abundance of methylesterified OG (GalAxMemAcn, x, m ≥ 1, n ≥ 0) or acetylated OG (GalAxMemAcn, x, m ≥ 0, n ≥ 1) from the emergence zone. **f**, Schematic representation of OG profiling experiment results. Top: control condition. Bottom: IAA-treated condition. Green and red arrows represent the CR tip and emergence zones, respectively. Plots representing the DP, DM and DA status of oligogalacturonides during the time course are displayed above each relevant zone. DP, degree of polymerization; DM, degree of methylesterification; DA, degree of acetylation.

### Homogalacturonan demethylesterification is homogeneously triggered by auxin in the rootlet emergence zone

The spatial regulation of pectin remodeling enzyme activity is known to be essential for organ growth and development (Levesque-Tremblay *et al.*, 2015). In cluster roots, auxin caused demethylesterification of HG in the tip zone and in the emergence zone (Fig. 3d, e and f). To understand whether the observed changes might be tissue or cell specific, we assessed the pattern of methylesterification of pectins in the emergence zone using a subset of monoclonal antibodies targeting HG of various DM status (Fig. 4). Immunolabeling of un-esterified HG with the LM19 antibody was uniform across rootlet primordium developmental stages and was not affected by auxin treatment (Fig. 4a). In contrast, JIM5, labeling HG stretches with low DM, displayed a strongly increased signal in cortex cells when treated with auxin (Fig. 4b). The area directly touching the rootlet primordium was heavily marked, which could be explained by the high number of walls from collapsed cells in this particular location. We also noticed that phloem cells directly below the pericycle cell layer were labeled with JIM5, but this tended to decline in auxin-treated roots. These data suggest that auxin reduces the degree of HG methylesterification. However, pectins seemed to be preponderantly present in fully demethylesterified forms in this zone (Fig. 3e). When looking specifically at highly methylesterified HG using LM20, we observed that the cortex cell three-way junctions were strongly labeled (Fig. 4c). In contrast, weak to no labeling was found in other cell types in the epidermis and the stele. Auxin substantially decreased LM20 labeling homogeneously in all cortical cells. In agreement, JIM7, which recognizes a highly methylesterified HG epitope, showed the same pattern as LM20 with a weaker signal in auxin-treated cluster roots (Fig. 4d). Overall, our results showed a clear positive effect of auxin in the removal of methylester groups from HG regardless of the location (overlaying a primordium or not) of the cortical cells.

**Figure 4:**
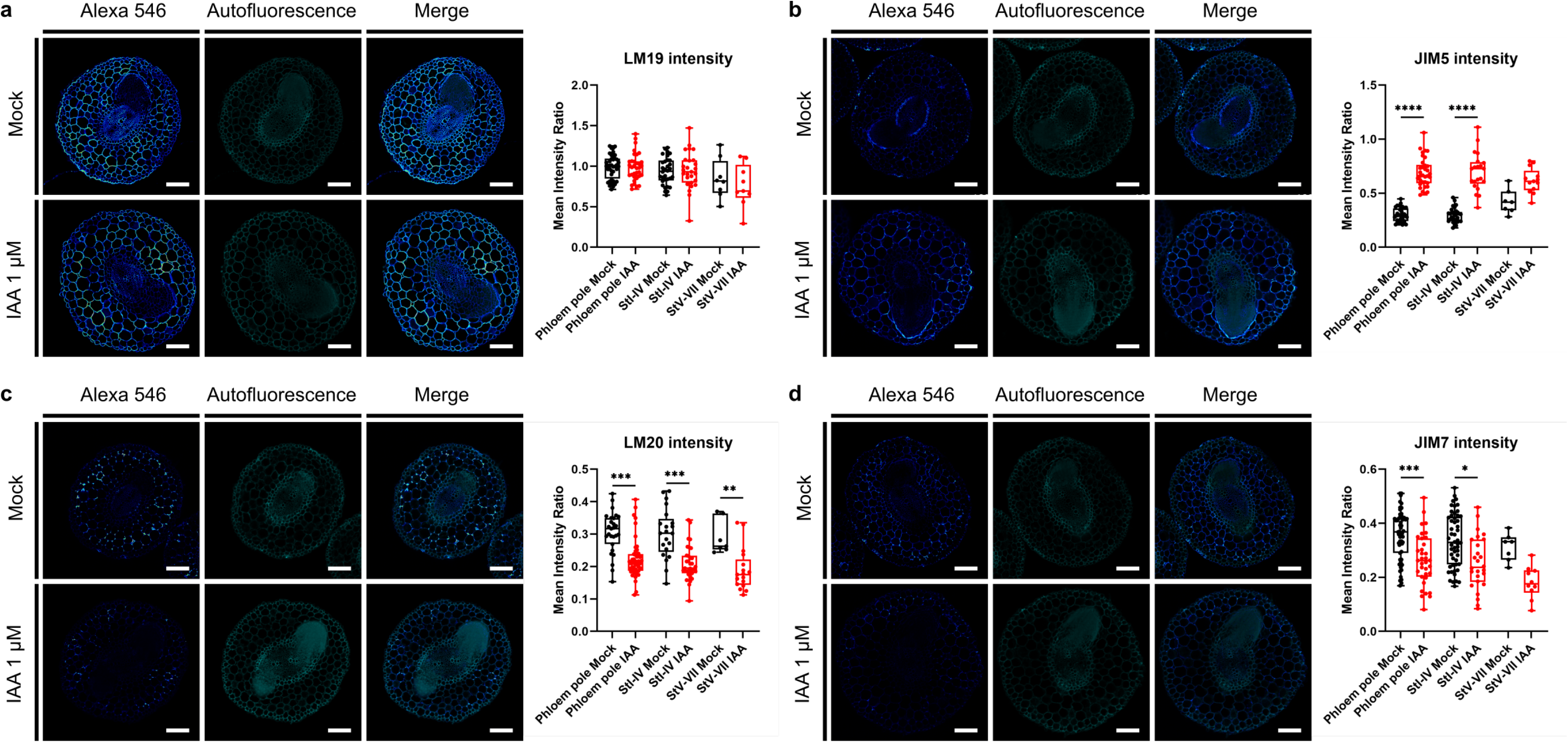
Auxin induces HG demethylesterification in cluster root cortical tissues independently of rootlet primordium emergence stage and location. Immunolabeling of the homogalacturonan specific LM19 (**a**), JIM5 (**b**), LM20 (**c**) and JIM7 (**d**) antibodies of cluster root cross-sections 2 days after treatment with 1 μM IAA or Mock (Ethanol 0.01%). Scale bar: 100 μm. Boxplot represents antibody intensity defined as the ratio of secondary antibody (Alexa 546) signal to autofluorescence signal in the "mechanically unchallenged" cortical tissues (phloem pole) or in that overlaying early (stage I to IV) and late (stage V to VII) rootlet primordia. All data points are displayed, whiskers show minimal and maximal values (**a**, mock: phloem pole: n = 40 sections, StI-IV: n = 32, StV-VII: n = 8; IAA: phloem pole: n = 34, StI-IV: n = 27, StV-VII: n = 9; **b**, mock: phloem pole: n = 36, StI-IV: n = 29, StV-VII: n = 7; IAA: phloem pole: n = 34, StI-IV: n = 21, StV-VII: n = 13; **c**, mock: phloem pole: n = 27, StI-IV: n = 21, StV-VII: n = 7; IAA: phloem pole: n = 46, StI-IV: n = 33, StV-VII: n = 17; **d**, mock: phloem pole: n = 59, StI-IV: n = 52, StV-VII: n = 7; IAA: phloem pole: n = 35, StI-IV: n = 26, StV-VII: n = 10). Statistical significance was computed with the Kruskal-Wallis test. ****: pVal < 0.001, ***: pVal < 0.005, **: pVal < 0.01, *: pVal < 0.05.

### (1,4)-β-D-galactan and extensins/type-I arabinogalactans are differentially distributed in primordium-overlaying cell walls

Rhamnogalacturonans-I (RG-I) are the second most abundant polymers in pectins and are composed of a backbone of alternating rhamnose and galacturonic acid with side chains of α-(1,5)-l-arabinans, β-(1,4)-galactans, and type-I arabinogalactans (Caffall et Mohnen, 2009). There is growing evidence that (1,4)-β-D-galactans are involved in cell-cell adhesion in many developmental contexts (Ng *et al.*, 2015, Moore *et al.*, 2014). To determine whether (1,4)-β-D-galactans could play a role in rootlet emergence we examined their distribution using the specific LM5 antibody in cluster root cross sections (Fig. 5). We noticed a significant reduction in the fluorescent signal for LM5 in the primordium-overlaying cortical cells from stage IV onward. Interestingly, epidermis inner and shared cell walls were not labeled indicating core cell wall differences between epidermis and cortex tissues. In contrast with HG methylesterification degree, RG-I (1,4)-β-D-galactan reduction was not altered by auxin treatment (Fig. 5). Contrarily to the clear depletion of LM5 signal from stage IV onward in the control, we observed an increased distribution of a subset of antibodies targeting putative sugar epitopes from cell wall glycoproteins (Extended Data Fig. 8). Extensin-specific JIM11 (Smallwood *et al.*, 1994) labeling was significatively increased in primordium-overlaying cortical cells in latter stages (Extended Data Fig. 8a). The same spatial distribution was observed for JIM93 and JIM94 antibodies for which the epitope is suggested to be found in the arabinogalactan side chain of hydroxyproline-rich glycoproteins (HRGP) (Pattathil *et al.*, 2010, Hall *et al.*, 2013) (Extended Data Fig. 8b, c). Labeling of JIM11, JIM93 and JIM94 were highly comparable in the stele, with an unusual labeling of the protoxylem, and in the walls separating the pericycle and endodermis cell layers. These results suggest extensive RG-I remodeling in cortical-overlaying cells that are mechanically challenged by the primordium outgrowth.

**Figure 5:**
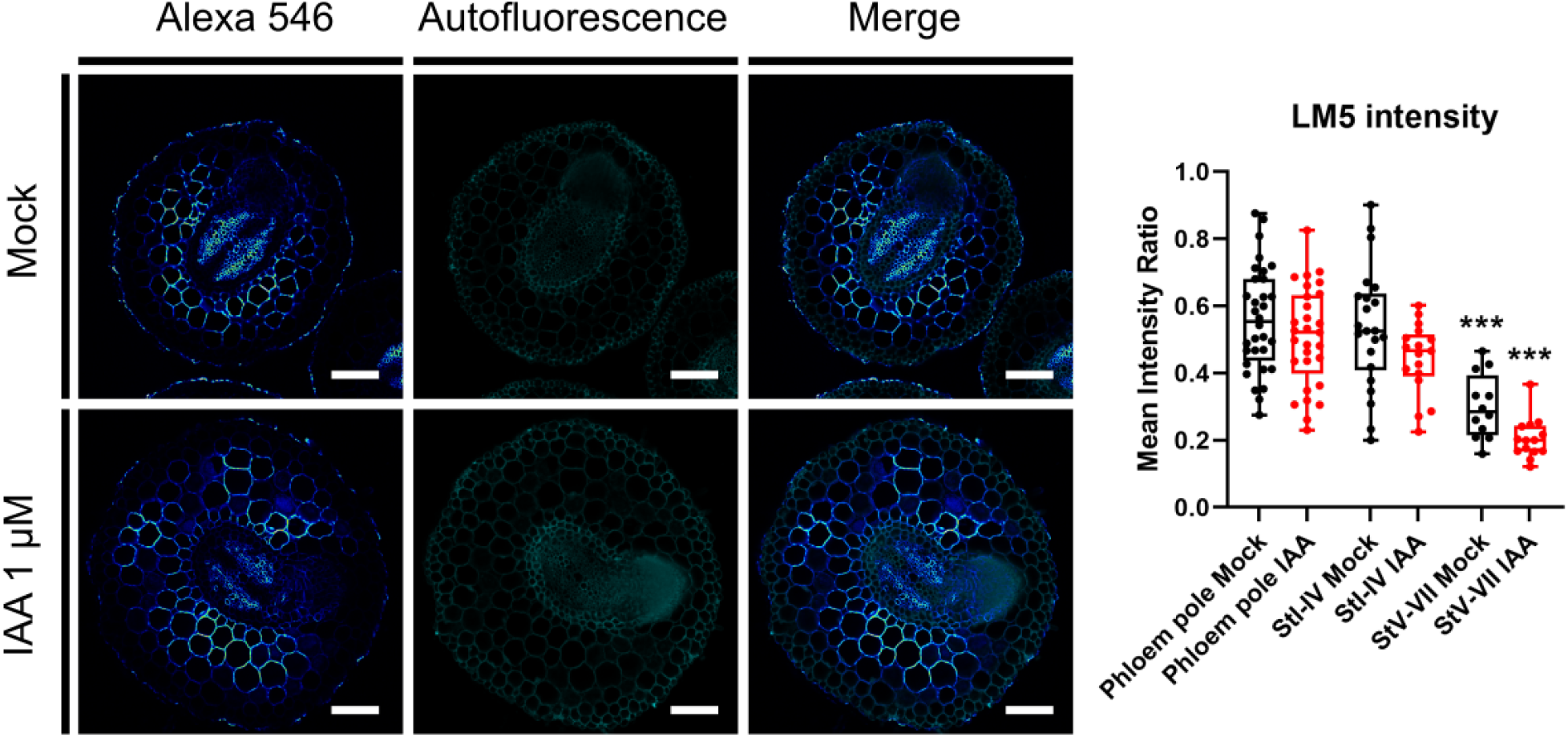
Rhamnogalacturonan I (1,4)-β-D-galactan is depleted in rootlet primordium-overlaying cells independently of auxin treatment. Immunohistochemistry with (1,4)-β-D-galactan specific LM5 antibody in cluster root cross-sections 2 days after treatment with 1 μM IAA or Mock (Ethanol 0.01%). Boxplot represents the antibody intensity defined as the ratio of secondary antibody (Alexa 546) signal to autofluorescence signal in the "mechanically unchallenged" cortical tissues (phloem pole) or those overlaying early (stage I to IV) and late (stage V to VII) rootlet primordia. All points are displayed, whiskers show minimal and maximal values (mock: phloem pole: n = 32 sections, StI-IV: n = 22, StV-VII: n = 12; IAA: phloem pole: n = 29, StI-IV: n = 17, StV-VII: n = 14). Statistical significance (compared to "phloem pole" mean intensity ratio mock conditions) was computed with the Kruskal-Wallis test. ***: pVal < 0.005. Scale bars: 100 μm.

## Discussion

Since more than a century, botanists have been arguing about whether the secondary root emerges purely mechanically or is helped by the digestion of the outer cortical layers or both (Pond, 1908). Using white lupin, Pond elegantly compared the lateral root primordium to a boat cutting through the water, making its way out by pushing away the outer cortical tissues. His microscopic observations determined that separation but not digestion of the outer cortical cells occurs in the late stages, corresponding to stage V onwards (Gallardo *et al.*, 2018). However, previous reports claimed that the lateral root secretes a substance actively digesting the outer cortical cells (Vonhöne, 1880), or that the passage of the primordium may be aided by enzymatic activity of the external tissue (Pfeffer, 1893). Both hypotheses have been put in the spotlight by the discovery in *Arabidopsis*, one century later, of the central action of primordium tip-derived auxin in the activation of cell wall remodeling enzyme genes in the overlaying cells (Swarup *et al.*, 2008). Our observation of cluster roots treated with auxin and with auxin transport inhibitors confirmed a positive role of the phytohormone on rootlet initiation and emergence (Fig. 1). IAA treatment modulated the expression of many genes in the cluster root emergence zone (Fig. 2b, c; Extended data Fig. 2; Supplemental Table 2). As the first outcome of auxin stimulus, several genes related to transcriptional regulation were upregulated such as *Lalb_Chr01g0019701* and *Lalb_Chr02g0142291*, the two closest orthologues of *LATERAL ORGAN BOUNDARIES-DOMAIN 29* (*LBD29*) in *Arabidopsis*, a transcriptional activator transducing auxin signals in outer cell layers (Porco *et al.*, 2016), suggesting that early developmental pathways are conserved across species.

We also observed in a second transcriptional wave caused by auxin, involving a steady upregulation of cell wall related genes (Fig. 2c, cluster 3), which is a response also found during lateral root development in *Arabidopsis* (Lewis *et al.*, 2013). We showed that in white lupin, an auxin-induced polygalacturonase, named *LaPG3*, is specifically expressed at the location of rootlet primordium emergence, in the outer cortex cell layers (Fig. 2d, Extended Data Fig. 7a). This enzyme shows structural similarities with fungal endo-polygalacturonase (Extended Data Fig. 6) and is the closest lupin orthologue of *AtPGLR* (*At5g14650*, Extended Data Fig. 3a), an acidic polygalacturonase identified as a putative candidate involved in lateral root emergence in *Arabidopsis* (Kumpf *et al.*, 2013, Hocq *et al.*, 2020). Although belonging to a multigenic family (108 members in *L. albus* according to Hufnagel *et al.*, 2020 genome release), amiRNA driven down-regulation of *LaPG3* in lupin hairy roots slightly delayed rootlet emergence (Fig. 2e). A simple explanation for this phenotype would be a reinforced cell wall and stronger cell adhesion in primordium-overlaying cells, slowing down the overall process of rootlet emergence. We also noticed the increased expression of auxin responsive *GH3* genes in *proPG3::amiR-PG3* cluster roots (Extended Data Fig. 7d). Auxin sensitivity is dampened in tobacco plants expressing fungal endo-PG and treatment with oligogalacturonide elicitors leads to the same effect in *Arabidopsis* (Ferrari *et al.*, 2008, Savatin *et al.*, 2011). Accordingly, we noticed a similar antagonism in the context of our microRNA targeting *LaPG3* transcript. Indeed, a low *LaPG3* expression could reduce oligogalacturonide-eliciting responses, thus alleviating the inhibition of auxin responses. This negative feedback loop between PG and auxin responses could help stabilize the rootlet emergence process by the inactivation of IAA via auxin-amido synthetase GH3.

Polygalacturonase activity requires structural modifications of the homogalacturonan pectic backbone and polygalacturonases are mostly active on demethylesterified stretches. Auxin plays an important role in the regulation of pectin methylesterase (PME) and PME inhibitor (PMEI) expression, which in turn modulates the degree of methylesterification in diverse organs. However, auxin action on pectin methylesterification depends on the developmental process at stake, positively regulating the removal of methylester groups in the shoot apical meristem while contrastingly promoting methylesterification during apical hook development (Braybrook et Peaucelle, 2013, Jonsson *et al.*, 2021). We were able to observe the dynamics of homogalacturonan methylesterification in the developing cluster root temporally, during early cluster root development, and spatially, discerning the most distal tip part (meristematic and elongation zone) and the rootlet emergence part (Fig. 3). Auxin triggered demethylesterification in the whole cluster root with the homogalacturonans in the tip zone being substantially more methylesterified than in the emergence zone (Fig. 3d, e, Fig. 4). A low degree of methylesterification was correlated with the release of galacturonic acids with a low degree of polymerization after *Aspergillus aculeatus* endo-PG processing (Fig. 3b – e, Extended data Fig. 7). This could be explained by the preferred affinity of AaPGM2 for demethylesterified galacturonic acid stretches (Safran *et al.*, 2021). Based on its predicted structure and high similarity with fungal endo-PG and *Arabidopsis* PGLR (Pickersgill *et al.*, 1998, Van Santen *et al.*, 1999, Cho *et al.*, 2001, Extended data Fig. 6), LaPG3 is likely to have a similar mode of action, however our attempts to produce recombinant enzyme for further characterization have failed so far. We hypothesize here that auxin causes pre-processing of the cell walls by regulating enzymes such as PME, allowing cell wall degradation in the emergence zone at specific locations marked by the expression of LaPG3. Such a tandem regulation of pectin degradation has been observed during pollen tetrad separation in *Arabidopsis*. *QUARTET1* (*QRT1*) encodes a PME that removes methylester groups, which stimulates *QUARTET3* (*QRT3*), a PG, to cleave homogalacturonans, allowing cell wall loosening (Rhee *et al.*, 2003, Francis *et al.*, 2006). *QRT1* is also specifically expressed at other locations of cell separation in *Arabidopsis*, including lateral root emergence-overlaying cells (Francis et *al.*, 2006). An analogous mechanism has been proposed for fruit softening in avocado and tomato (Wakabayashi *et al.*, 2003).

The outer cortex is subject to enormous mechanical constraint in the rootlet emergence zone due to the growing primordium. Cell death can be observed in the endodermis and to a lesser extent in the cortex cells in species such as *Arabidopsis* (Escamez *et al.*, 2020). In white lupin, those layers (endodermis and inner cortex) are dividing and become included within the rootlet primordium. Microscopic observations of late-stage primordia (Stage V onwards) indeed show collapsed outer cortex cells but whether cell death occurs remains to be proven (Gallardo *et al.*, 2018). Once the most difficult part of the rootlet’s journey has been dealt with by the recruitment of endodermis and inner cortical cell divisions, one can assume that late emergence solely takes place due to the displacement of outer cortex and epidermis cells. The specific expression of LaPG3 in those cells when a primordium grows underneath suggests however that degradation of the pectin-rich middle lamellae is crucial to reduce cell adhesion. Still, LaPG3 expression was not found in epidermis suggesting that cell separation might occur through a different mechanism. This could be explained by a different cell wall composition. Immunolabeling experiments with antibodies targeting homogalacturonan epitopes with diverse degrees of methylesterification confirmed that auxin triggered demethylesterification in the emergence zone. However, we were not able to identify differential distribution of methylesterified and demethylesterified homogalacturonans in cortex cell walls challenged by a primordium or not (Fig. 4). Such a differential distribution of methylesterified pectins was recently reported at the junction between the lateral root primordium and endodermis in the early stages and later between endodermis and cortex in *Arabidopsis* (Waschman *et al.*, 2020). Our observations of homogalactan epitopes were conducted at a late stage of primordium emergence in a region of the cluster root that experiences hundreds of primordia growing in close proximity. The most striking observations of differential cortex cell wall composition were revealed using antibodies that recognize rhamnogalacturonan-I and glycoprotein epitopes (Fig. 5, Extended Data Fig. 8). Rhamnogalacturonans-I are the second most represented component of the pectin matrix. The complex rhamnogalacturonan-I main chain is composed of α-D-galacturonosyl and rhamnosyl residues, with side chains of (1,4)-β-D-galactan and (1,5)-α-L-arabinan residues of various degrees of polymerization. This pectin component is thought to play a role in cell adhesion through crosslinks with cellulose microfibrils (Lin *et al.*, 2015). Notably, a loss of (1,4)-β-D-galactans has been correlated with fruit softening in the ripening process of apple and grapes (Ng. *et al.*, 2015, Moore *et al.*, 2014). It is also worth mentioning that (1,4)-β-D-galactan side chains contribute to the gelation capacity of rhamnogalacturonan-I *in vitro* (Mikshina *et al.*, 2017). In this study, we detected a loss of (1,4)-β-D-galactan epitopes in the primordium-overlaying cortical cells, for which the effect was independent of auxin (Fig. 5). The epidermis inner cell wall and the direct subepidermal cell layer were intriguingly not marked, reflecting different pectin composition and possibly mechanical properties of those cells. Nevertheless, it remains to be determined whether (1,4)-β-D-galactan loss in rhamnogalacturonans-I is part of a controlled active process of cell separation or is a passive consequence of cell wall remodeling.

The same question needs to be investigated for the visible increase of extensin and HRGP epitope labeling at the location of primordium emergence (Extended Data Fig. 8). Extensins are proteins known to be involved in cell wall relaxation especially in rapidly growing pollen tubes and root hairs (Velazquez *et al.*, 2011, Wang *et al.*, 2018) but their role in cell adhesion is more enigmatic. Nonetheless, a high expression of an extensin coding gene has been correlated to tomato fruit ripening, thereby possibly affecting cell wall cohesion (Ding *et al.*, 2020). JIM11, JIM93 and JIM94 labeled the same tissues in the cluster root emergence zone, but the epitopes are not fully characterized. However, their distribution suggests binding to a highly similar glycan epitope (Pattathil *et al.*, 2010). The strong labeling of JIM11, JIM93 and JIM94 could be due to two possibilities. Firstly, an accumulation of extensins and HRGP in the challenged cortical cell wall. Their accumulation in primordium-overlaying cells could lead to cell wall softening and cell separation. Secondly, possible pectin remodeling, suggested here by auxin induced demethylesterification and spatial PG expression during rootlet emergence in addition to the depletion of (1,4)-β-D-galactan residues, could increase the epitope availability. Thus, more extensin and HRGP antibody signal could be caused by increased abundance of the protein and/or by uncovering epitope sites in modified cell walls. This is likely to occur after the auxin-induced changes leading specifically to homogalacturonan modifications (demethylesterification) and degradation (polygalacturonase activity). However, it cannot be excluded that a solely mechanical aspect could trigger those cell wall modifications.

## Conclusion

In this study, we used white lupin (*Lupinus albus*) as a model to understand the contribution of auxin and cell wall remodeling during rootlet emergence because of its striking secondary root developmental phenotype. Auxin accelerates primordium emergence and causes a shift in the transcriptional landscape including the upregulation of cell wall related genes. Among several candidate genes, we identified *LaPG3*, an endo-polygalacturonase gene expressed in the outer cortex and positively regulating rootlet emergence. Additionally, we found that homogalacturonans, the substrate of polygalacturonase, were highly demethylesterified in the emergence zone, an effect amplified by external auxin treatment. In contrast to methylesterified homogalacturonans, we noticed that (1,4)-β-D-galactans from rhamnogalacturonan-I side chains were differentially distributed in mechanically challenged cortex cell wall by the growing primordium. Conversely, we found that extensins and HRGP epitopes were enriched in the walls of those cells. Altogether, we propose a model for auxin-controlled cell separation during secondary root emergence in which auxin triggers homogalacturonan demethylesterification in the cluster root, rending possible the degradation of pectins in the cortex cells expressing *LaPG3*. The resulting cell wall loosening and loss of cell adhesion are accompanied by depletion of (1,4)-β-D-galactans and accumulation of extensins/arabinogalactan protein.

## Acknowledgements

We acknowledge Carine Alcon for technical support at the imaging facility MRI (Montpellier, France), a member of the national infrastructure France-BioImaging infrastructure supported by the French National Research Agency (ANR-10-INBS-04, «Investments for the future») and the microscopy facility at UPSC (Umeå, Sweden). We thank Serge Pilard (Analytical Platfrom, UPJV, France) for the LC-MS/MS analyses. This work was supported by the Kempestiftelserna (Scholarship SMK 1759, F.J.), the Swedish Research Council Vetenskapsrådet grant VR-2020-03420 (F.J.), the Knut and Alice Wallenberg Foundation and Vinnova (Verket för Innovationssystem) (F.J., S.R.). This project has received funding from the European Research Council (ERC) under the European Union’s Horizon 2020 research and innovation program (Starting Grant LUPINROOTS - grant agreement No 637420 to B.P).

## List of Extended Data

Extended Data Fig.1: Representative pictures of nine-day-old *Lupinus albus* grown in hydroponic medium after two days supplemented with 0.01% ethanol (control treatment – two plants), 10 μM NPA, 1 μM IAA, 2.5 μM 1-NOA or 25 μM 1-NOA. Scale bar: 5 cm.

Extended Data Fig.2: Minor clusters identified in the transcriptome of auxin treated cluster roots.

Expression pattern of minor clusters identified in the main Figure 2b (light blue, purple, black and green bars on the heatmap). Bubble plots display gene ontology enrichment analysis with the identified molecular (red), biological process (green) and cellular component (blue) associated with the differentially expressed genes among the clusters.

Extended Data Fig.3: Identification of white lupin cell wall related putative orthologues and promoter activity in transformed hairy root.

**a**, Phylogenetic tree of cell wall related genes based on protein sequences in *L. albus* (green dot) and *Arabidopsis*. The more distantly related transmembrane amino acid transporter protein GAMMA-AMINOBUTYRIC ACID TRANSPORTER 1 (GAT1, *At1g08230*) was chosen as an outgroup for the analysis. **b**, Representative promoter::GUS pictures for each putative *L. albus* cell wall related gene in cluster root tips and rootlet primordia in the different stages of development described by Gallardo *et al.*, 2018. Cross sections in the emergence zone are also shown. Scale bar: 100 μm.

Extended Data Fig.4: Promoter activity of white lupin auxin responsive cell wall genes in transformed hairy root.

Representative promoter::GUS pictures for *L. albus* auxin responsive cell wall related genes in cluster root tips and in rootlet primordia in the different stages of development described by Gallardo *et al.*, 2018. Cross sections in the emergence zone are also shown. Scale bar: 100 μm. Asterisk shows the cross section for *pLaEXP2::GUS* in the elongation zone of the cluster root (not in the emergence zone), where gene expression was found. Expression level of the candidate genes over the course of the auxin transcriptome generated in this study (see Fig. 2) are shown on the right.

Extended Data Fig.5: Auxin-responsive *Lupinus albus POLYGALACTURONASE3* (*LaPG3*) expression.

RNAseq expression level of *LaPG3*, an auxin-responsive gene from the expression cluster 3.

Extended Data Fig.6: LaPOLYGALACTURONASE3 *in silico* characterization.

**a**, Protein alignment of AtPGLR (*AT5g14650*) and LaPG3 (*Lalb_Chr02g0160451*) using clustal omega (https://www.ebi.ac.uk/Tools/msa/clustalo/. version 1.2.4). Results are shown using MView, **b**, Surface and ribbon representation of LaPG3 modeled structure (gray) with amino acid of the active site highlighted (blue). **c**, LaPG3 modelled structure (grey) superimposed to the structure of *Pectobacterium carotovorum* PG (PDB:1BHE, yellow), *Aspergillus niger* PGII (PDB:1CZF, green) and *Aspergillus acuelatus* PG (PDB:1IA5, purple). **d**, Amino acids of the LaPG3 active site (blue) superimposed onto that of the *Aspergillus acuelatus* active site (yellow).

Extended Data Fig.7: Characterization of *LaPOLYGALACTURONASE3* microRNA lines in white lupin hairy roots.

**a**, Longitudinal view of proLaPG3::nlsYFP localization in cluster roots during rootlet primordium emergence, stages III-VI. proUBI-mCherry is a transformation internal positive control. White dashed lines outline rootlet primordia. Scale bars: 50 μm. **b**, Alignment of the artificial microRNA targeting its corresponding *LaPG3* transcript region (*amiR-PG3*). **c**, Rootlet number, cluster root length and rootlet density in hairy root composite plants expressing *proPG3::amiR-PG3* (n = 27 transformed roots) and *proPG3::nlsYFP* (n = 28) as a control. Statistical significance was computed using an unpaired t-test: ns: non-significant. **d**, Relative gene expression of *LaPG3*, *LaGH3.3*, *LaGH3.5* and *LaGH3.6* in *proPG3::amiR-PG3* transformed cluster roots compared to *proPG3::nlsYFP*. Gene expression values are relative to the expression in *proPG3::nlsYFP*, for which the value is set to 1. Error bars indicate SEM obtained from three independent biological replicates. Statistical significance was computed using an unpaired t-test: *: pVal < 0.05, **: pVal < 0.01, ***: pVal < 0.005.

Extended Data Fig.8: JIM11 and JIM93/JIM94 epitope labeling is increased in rootlet primordium-overlaying cells at late stages.

**a**, Immunolabeling of the extensin specific JIM11 antibody of cluster root cross-sections. **b**, Immunolabeling of the type I arabinogalactan chain putative JIM93 antibody of cluster root cross-sections. **c**, Immunolabeling of the type I arabinogalactan chain putative JIM94 antibody of cluster root cross-sections. Boxplots represent antibody intensity defined as the ratio of secondary antibody (Alexa 546) signal to autofluorescence signal in the "mechanically unchallenged" cortical tissues (phloem pole) or those overlaying early (stage I to IV) and late (stage V to VII) rootlet primordia. All points are displayed, whiskers show minimal and maximal values (**a**, phloem pole: n = 32, StI-IV: n = 16, StV-VII: n = 16; **b**, phloem pole: n = 29, StI-IV: n = 20, StV-VII: n = 11; **c**, phloem pole: n = 19, StI-IV: n = 11, StV-VII: n = 8). Statistical significance (compared to "phloem pole" mean intensity ratio) was computed with the Kruskal-Wallis test. ****: pVal < 0.001, **: pVal < 0.01. Scale bars: 100 μm.

## List of Supplemental Tables

Supplemental Table 1: DEG counts from RNAseq

Supplemental Table 2: GO enrichment analysis

Supplemental Table 3: Primers list

Supplemental Table 4: Antibodies list

## Materials and Methods

### Plant materials and growth conditions

White lupin (*Lupinus albus*) cv. AMIGA (from Florimond Desprez, France) used in this study were germinated in vermiculite for and grown three days under long day conditions (16 h light/8 h dark, 25 °C day/20 °C night, 65% relative humidity and PAR intensity 200 μmol.m².s⁻¹). The three-day-old seedlings were transferred to 1.6 L pots containing a phosphate-free hydroponic solution composed of the following: 400 μM Ca(NO_3_); 200 μM K_2_SO_4_; 54 μM MgSO_4_; 0.24 μM MnSO_4_; 0.1 μM ZnSO_4_; 0.018 μM CuSO_4_; 2.4 μM H_3_BO_3_; 0.03 μM Na_2_MoO_4_; 10 μM Fe-EDTA. Hydroponic medium was renewed weekly and permanent oxygenation was provided by an air pump.

### Chemical treatments and cluster root phenotyping

All chemicals used in this study were applied seven days after germination, at the onset of rootlet primordium development, directly in the hydroponic medium at the desired concentrations. The four upper cluster roots were sampled 48 hours after treatments and cleared in saturated aqueous chloral hydrate solution (250 g in 100 mL water) for two weeks before observation. Cluster root length, rootlet density and developmental stages of primordia were scored using a color camera on Olympus BX61 epifluorescence microscope (Tokyo, Japan). For artificial microRNA cluster root characterization, we sampled individual cluster roots (each is an independent transformation event) of 4 to 6 hairy root composite plants one week after hydroponic transfer and screened them under the confocal microscope (Zeiss LSM780) searching for ubiquitous mCherry signal at 561 nm (internal control of transformation). Hairy root experiments were conducted twice. Transformed cluster roots were transferred to chloral hydrate solution for phenotyping or frozen in liquid nitrogen for RNA extraction experiments. Transformed cluster roots showing obvious developmental irregularities such as root fusion or supernumerary xylem poles were excluded from the analysis.

### Gene expression analysis

Total RNA was extracted from 150 mg of cluster roots (3 to 4 roots) using the RNeasy Plant Mini Kit (Qiagen, 74904) and treated with the DNA-free DNA Removal Kit (Thermo Fisher Scientific, AM1906). 1 μg of total RNA was used for reverse transcription with the Invitrogen SuperScript III Reverse Transcriptase (Thermo Fisher Scientific, 18080093). Quantitative PCR was performed with the SsoAdvanced Universal SYBR Green Supermix (BioRad, 1725271) on a CFX Maestro 96 thermocycler (BioRad). Relative expression was calculated with the delta-delta Ct method, using the validated *LaNORM1* reference gene. Primers are available in Supplementary Table 3.

### RNA sequencing

Four independent biological replicates were used to prepare RNA-sequencing libraries with Illumina TruSeq Stranded Total RNA Kit with Ribo-Zero. The libraries were sequenced as 150bp pair-end reads using Illumina HiSeq3000 at Get-PlaGe core facility (INRA, Toulouse, France). A total of 2,110,906,218 paired-end reads were generated. The following procedure was applied to each paired read dataset. Cutadapt (version 1.15) was used to remove Illumina Truseq adapters from the sequencing data and to remove bases with a quality score lower than 30, in both 5’ and 3’ end of the reads. Reads with a length lower than 60 were discarded. The quality checked RNA-seq reads were then used to quantify white lupin transcript abundance using salmon (version 0.12.0) with the options validateMappings, gcBias and seqBias turned on. Data were imported to DESeq2 using tximpot and normalized using DESeq2 method. Then normalized counts were extracted for further analyses. Differential expression was performed using the following method: given a gene, if the mean count of the less expressed condition was more than twice inferior to the mean count of the most expressed condition, and if the mean of the most expressed condition was superior to 100, the gene was considered differentially expressed (DE). The DE gene count data for relevant samples were “log regularized”, using the Variance Stabilizing Transformation (vst) function from DESeq2. Then the degPatterns function from the DEGReport package was used to group genes into clusters based on their expression profile using the divisive hierarchical clustering method diana from the cluster package. Gene Ontology (GO) annotation was then performed using the available GO annotation for white lupin that can be found in the first version of the white lupin genome annotation file (https://www.whitelupin.fr/download.html). The package goseq was used to find GO enrichment in each of the clusters previously computed. Only enriched GO terms associated with a pvalue < 0.01 were kept.

### Phylogenetic trees

Protein sequences from white lupin were retrieved from the white lupin genome database (www.whitelupin.fr – Hufnagel *et al.*, 2020). Sequences producing significant alignments after blastp analysis of full length protein sequences of *Arabidopsis* as queries (Total score > 700; Evalue==0) were considered as putative lupin orthologous proteins for each gene family. Sequences were aligned by MUSCLE and evolutionary history was inferred using the Neighbor-Joining method (bootstrap replication number: 1000). Phylogenetic trees were then exported from the software MEGA X (version 10.0.4).

### Promoter GUS cloning

Upstream genomic sequences of white lupin cell wall related genes (primers, gene name and promoter length are listed in Supplemental table 3) were amplified by PCR from *L. albus* cv. AMIGA genomic DNA. The amplified fragments were cloned using the Gateway BP clonase enzyme mix into the pDONR201 entry vector (Thermo Fisher Scientific, 11798013) transformed in *E. coli* TOP10, verified by sequencing and recombined into the binary vector pKGWFS7 containing a GFP-GUS fusion (Karimi *et al.*, 2002) using the Gateway LR clonase II enzyme mix (Thermo Fisher Scientific, 11791020).

### Artificial microRNA cloning

Artificial microRNA (amiR) targeting *LaPG3* transcripts was designed using the online tool WMD3 (http://wmd3.weigelworld.org/cgi-bin/webapp.cgi) and the pRS300 backbone vector according to the protocol of Schwab *et al.*, 2006. The amiR-LaPG3 and the nlsYFP coding sequence (transformation control) were inserted into the entry vector pDONR201 (P1P2 recombination sites) and the promoter of *LaPG3* was inserted into the pDONRP4P1r (sequences listed in Supplemental table 3). The two fragments were recombined into the multisite Gateway vector pK7m24GW_CR, specifically designed for hairy root transient expression and cluster root phenotype analysis in this study, using Gateway LR clonase II enzyme mix. The same procedure was repeated to generate the DR5::nlsYFP construct in pK7m24GW_CR.

### Hairy root transformation

*Rhizobium rhizogenes* strain *ARqua-1* was used for hairy root transformation of white lupin. The bacteria were transformed with the expression vector by electroporation. LB agar plates were supplemented with 100 μM acetosyringone and appropriate antibiotics and inoculated with 200 μl of liquid bacterial culture. The plates were incubated at 28 °C for 24 h, producing a fresh and dense bacterial lawn used for lupin seedling transformation. Meanwhile, white lupin seeds were surface sterilized 30 min in bleach (Halonet 20%) and washed four times in sterile water. Seed were germinated on half MS medium (pH 5.7). Two days after germination, radicles of 1 cm were cut at 0.5 cm from the tip with a sterile scalpel. The wounded part was inoculated with the *Rhizobium rhizogenes* lawn. Inoculated seedlings were placed on square agar plates (0.7 % agar in 1X Hoagland solution) containing 15 μg/mL Kanamycin. Hoagland medium without phosphate is composed of 200 μM MgSO_4_; 400 μM Ca(NO_3_)2; 325 μM KNO_3_; 100 μM NH_4_Cl; 10 μM Na-Fe-EDTA; 9.3 μM H_3_BO_3_; 1.8 μM MnCl2; 0.17 μM ZnSO_4_; 0.06 μM CuSO_4_; 2.3 μM Na_2_MoO_4_. The plates were placed in a growth chamber under long day conditions (16 h light/8 h dark, 25 °C day/20 °C night). After seven days, seedlings were transferred to vermiculite in a mini greenhouse. Ten days later, the plants growing hairy roots were transferred to hydroponics (see “Plant materials and growth conditions” section).

### GUS Histochemical assay

Cluster roots were harvested from transgenic plants and immediately fixed in ice-cold 90% acetone for 30 minutes, washed three times in 0.1 M phosphate buffer (pH 7) and incubated in X-Gluc buffer (0.1% X-Gluc; 50 mM phosphate buffer, pH 7, 2 mM potassium ferricyanide, 2 mM potassium ferrocyanide, 0.05% Triton X-100) for 1 to 24 hours depending on the construct. X-Gluc buffer was then removed and replaced by saturated aqueous chloral hydrate solution to allow clearing of the tissues.

### Microscopic analysis

GUS-stained, *DR5::nlsYFP* and *LaPG3::nlsYFP* transformed cluster roots were embedded in 4% agarose (m/v) and cut with a vibratome to produce thick sections of 70 μm, (VT1000S, Leica Microsystems). The root sections were mounted on slides in 50% glycerol. Wild type (cv. AMIGA) cluster root thin sections of 6 μm were produced using a microtome (RM2165, Leica Microsystems). They were counterstained for 5 min either with 0.05% toluidine blue or with 0.1% ruthenium red in 1X phosphate buffer saline (1X PBS pH 7.4, Sigma Aldrich, P3813). GUS-stained and wild type cluster root sections were observed with a color camera on an Olympus BX61 epifluorescence microscope (Tokyo, Japan) with Camera ProgRes®C5 Jenoptik and controlled by ProgRes Capture software (Jenoptik, Jena, Germany). *DR5::nlsYFP* and *LaPG3::nlsYFP* transformed cluster roots were observed with a confocal microscope (Zeiss LSM 780, details in “confocal microscopy” section below).

### Immunolabeling experiments

Cross sections of 70 μm obtained with the vibratome were transferred onto chamber slides (Lab-teak, 177402) for immunostaining. They were first rinsed in 0.1 M glycine in 1X PBS and then twice in 1X PBS, each for 10 min. Sections were immersed in blocking buffer containing 5% bovine fetal serum (Sigma Aldrich, A9418) in 1X PBS at 4 °C overnight under gentle agitation. Monoclonal primary antibodies (list in Supplemental table 4) diluted 1/10 in the blocking buffer, were applied overnight at 4 °C under mild agitation. The sections were washed 3 times in 1X PBS for 10 min followed by two hours incubation in the secondary antibody (list in Supplemental table 4) diluted 1/500 in blocking buffer under gentle agitation. Sections were then washed three times in 1X PBS under mild agitation, 10 min each. The chambers were removed, and sections mounted in 50% glycerol prior to observation. Immunolabeling experiments were repeated three times.

### Confocal microscopy

Immunostained root sections were imaged on a confocal microscope (Zeiss LSM 780). Autofluorescence observation was performed using an argon laser at 405 nm and secondary antibodies were excited at 561 nm. Both were detected at a 566-679 nm window using the same settings (gain, offset, resolution) to allow quantification measurements. For *LaPG3::nlsYFP*, *DR5::nlsYFP* and screening of transformed cluster roots, mCherry internal control and nlsYFP were excited at 561 nm and 514 nm respectively and detected at 583-696 nm and 519-583 nm windows, respectively. Observations were made using Plan-Apochromat 10x/0.45 M27 and 20x/0.8 M27 objectives. Image acquisition was performed with the Zeiss ZEN black 2010 software and image analysis was conducted using the ZEN blue 2.3 lite software (Carl Zeiss Microscopy).

### Oligogalacturonide characterization and quantification

OG characterization and quantification in cluster root samples from the emergence zone and tip zone were performed using the method published in Voxeur *et al.*, 2019.

### Homology modeling

The LaPG3 model was created using I-TASSER software for protein structure and function prediction (https://zhanglab.ccmb.med.umich.edu/I-TASSER/; Zhang, 2008) with *Aspergillus aculeatus* rhamnogalacturonase A (PDB: 1RMG, Petersen *et al.*, 1997) as the best template. While these enzymes share rather low sequence identity (24.93%), they have high structural homology with estimated RMSD 6.6±4.0Å. The LaPG3 model consisted of 425 AA excluding the 24 AA of the signal peptide. The modeled structure was compared with that of *Pectobacterium carotovorum* PG (PDB:1BHE), *Aspergillus niger* PGII (PDB:1CZF) and *Aspergillus aculeatus* PG (PDB:1IA5). UCSF Chimera (http://www.cgl.ucsf.edu/chimera/) was used for creation of graphics (Pettersen *et al.*, 2004).

### Statistical analysis

Statistical analysis for phenotyping experiments and confocal image analysis were performed using GraphPad Prism version 9.0.2 for Windows (GraphPad Software, San Diego, California USA).

## Data availability

White lupin gene identifiers and full genomic sequences are available on the White Lupin Genome Portal (Hufnagel *et al.*, 2020; www.whitelupin.fr). The RNAseq data have been deposited at NCBI under the temporary name “SUB9968787”, bioproject “SAMN20089781”.

