## Supplementary figures for "Auxin and pectin remodeling interplay during rootlet emergence in white lupin"

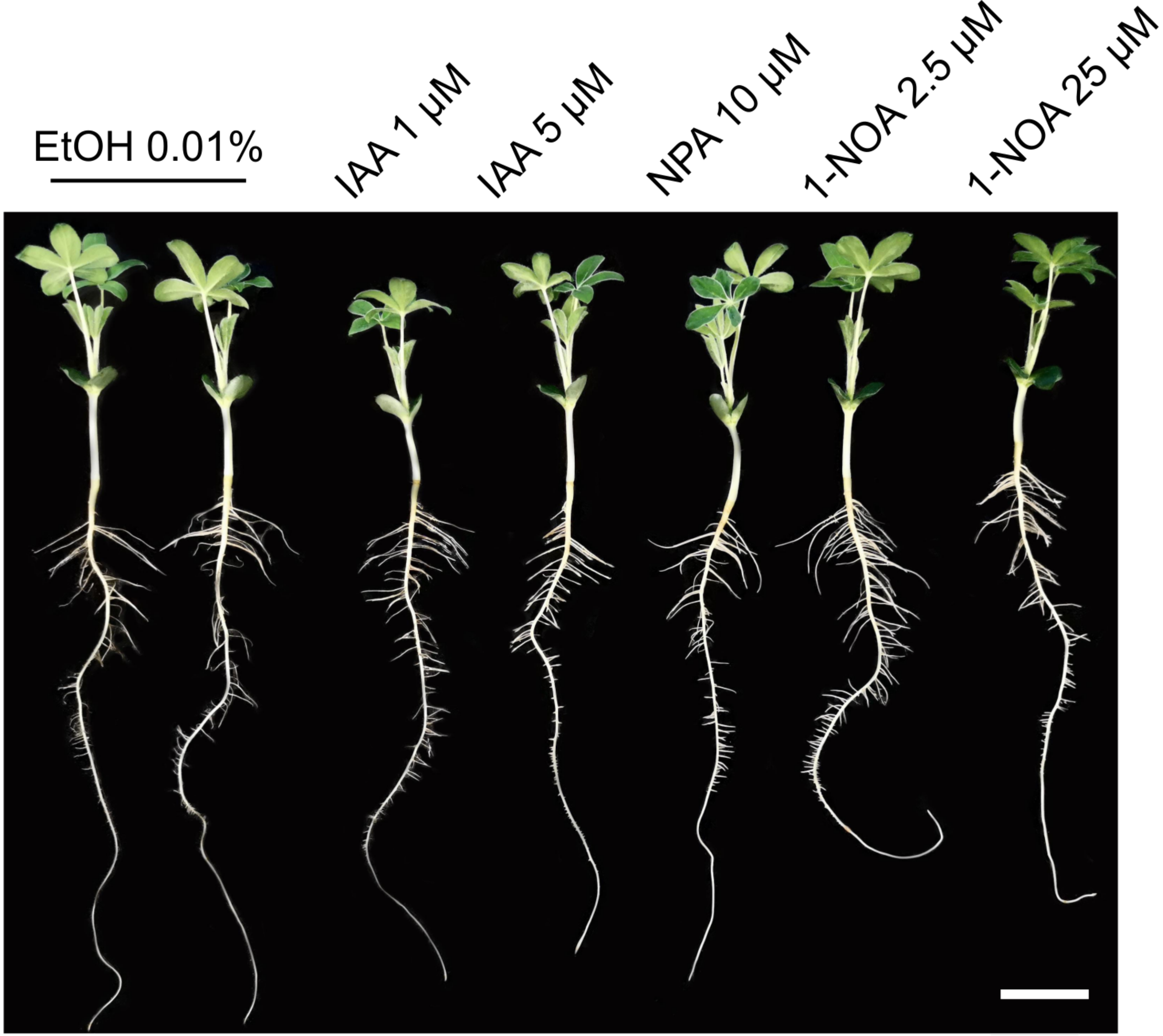


**Extended Data Fig.1: Representative pictures of nine days-old *Lupinus albus* grown in hydroponic medium after two days supplemented with 0.01% ethanol (control treatment - two plants), 10 µM NPA, 1 µM IAA, 5 µM IAA, 2.5 µM 1-NOA or 25 µM 1-NOA. Bar scale: 5 cm.**

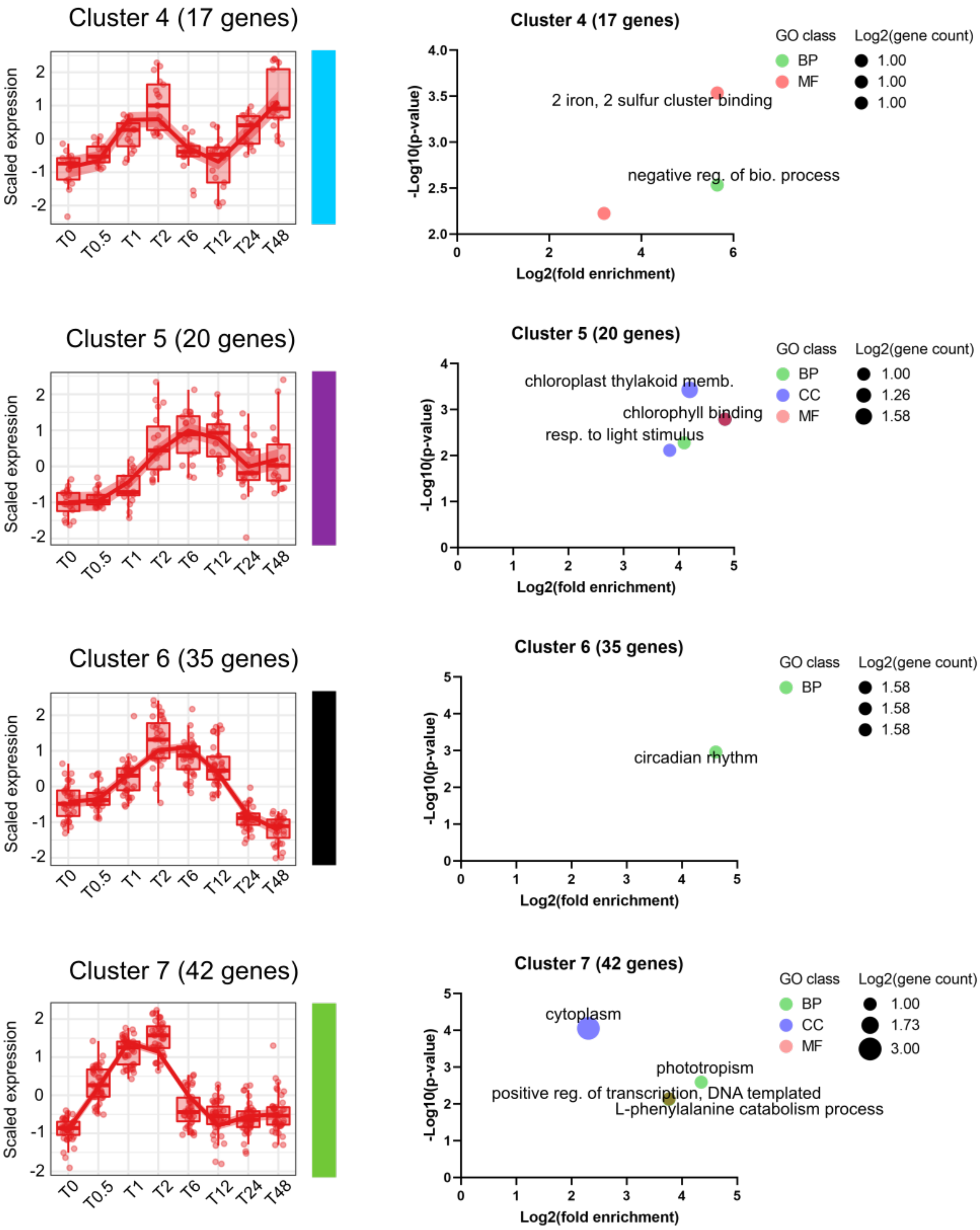


**Extended Data Fig.2: Minor clusters identified in the transcriptome of auxin treated cluster roots.**

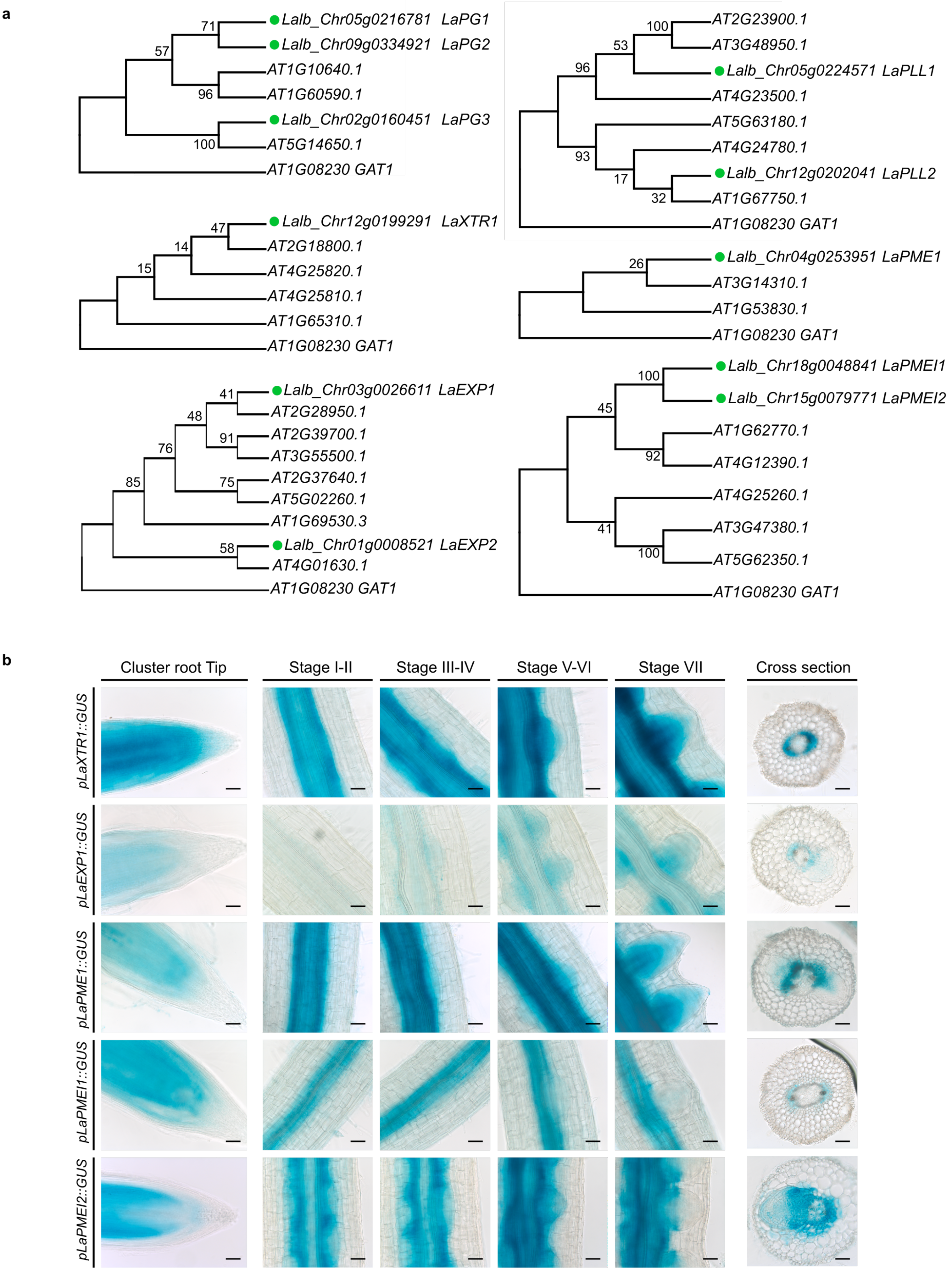


**Extended Data Fig.3: Identification of white lupin cell wall related putative orthologues and promoter activity in transformed hairy root.**

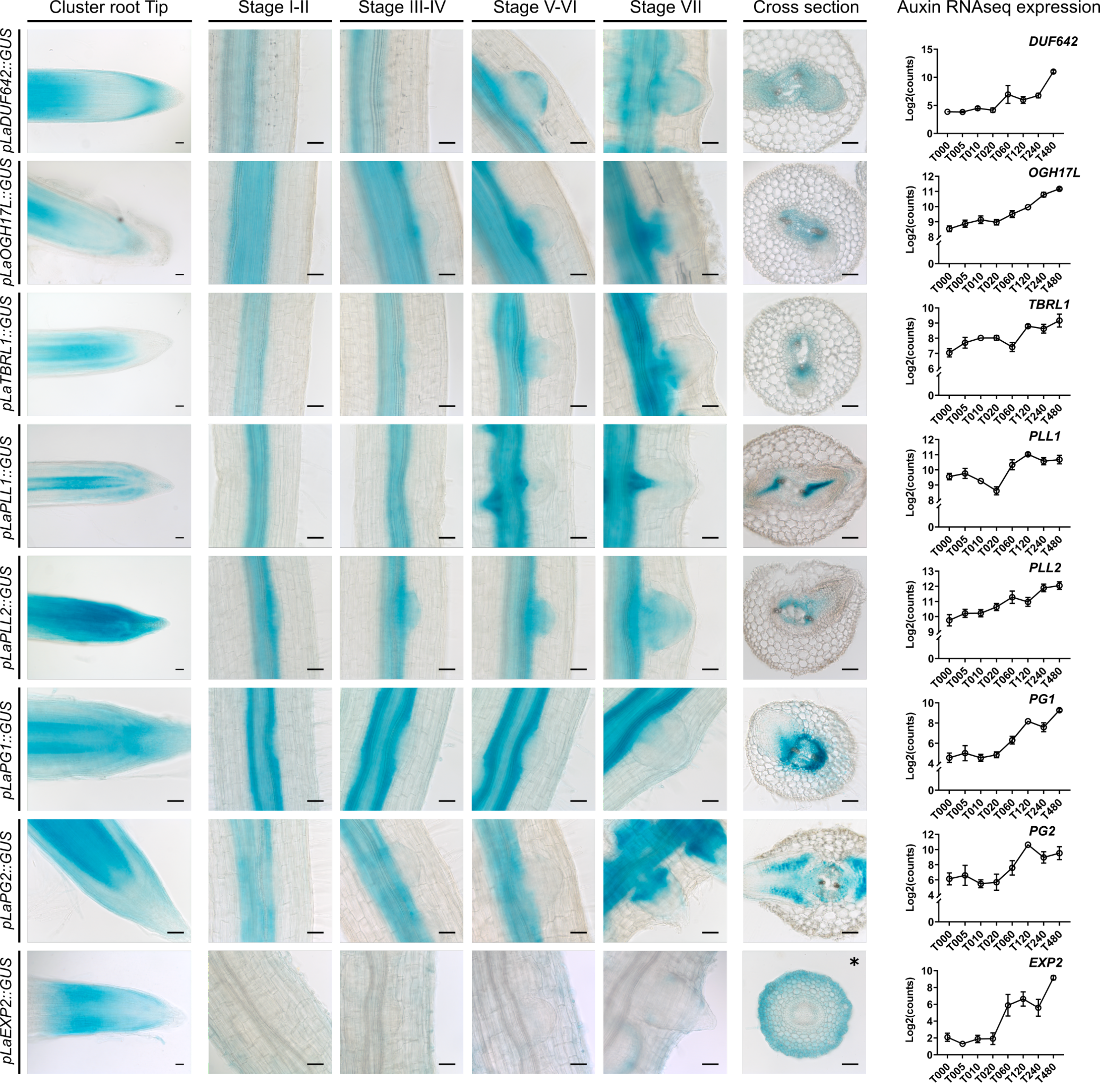


**Extended Data Fig.4: Promoter activity of white lupin auxin responsive cell wall genes in transformed hairy root.**

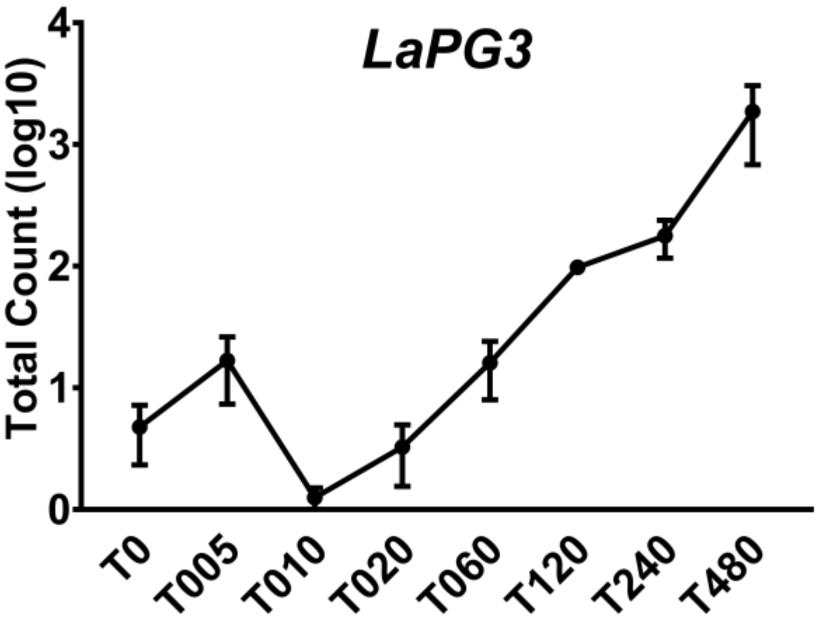


**Extended Data Fig.5: Auxin-responsive *Lupinus albus POLYGALACTURONASE3* (*LaPG3*) expression.**

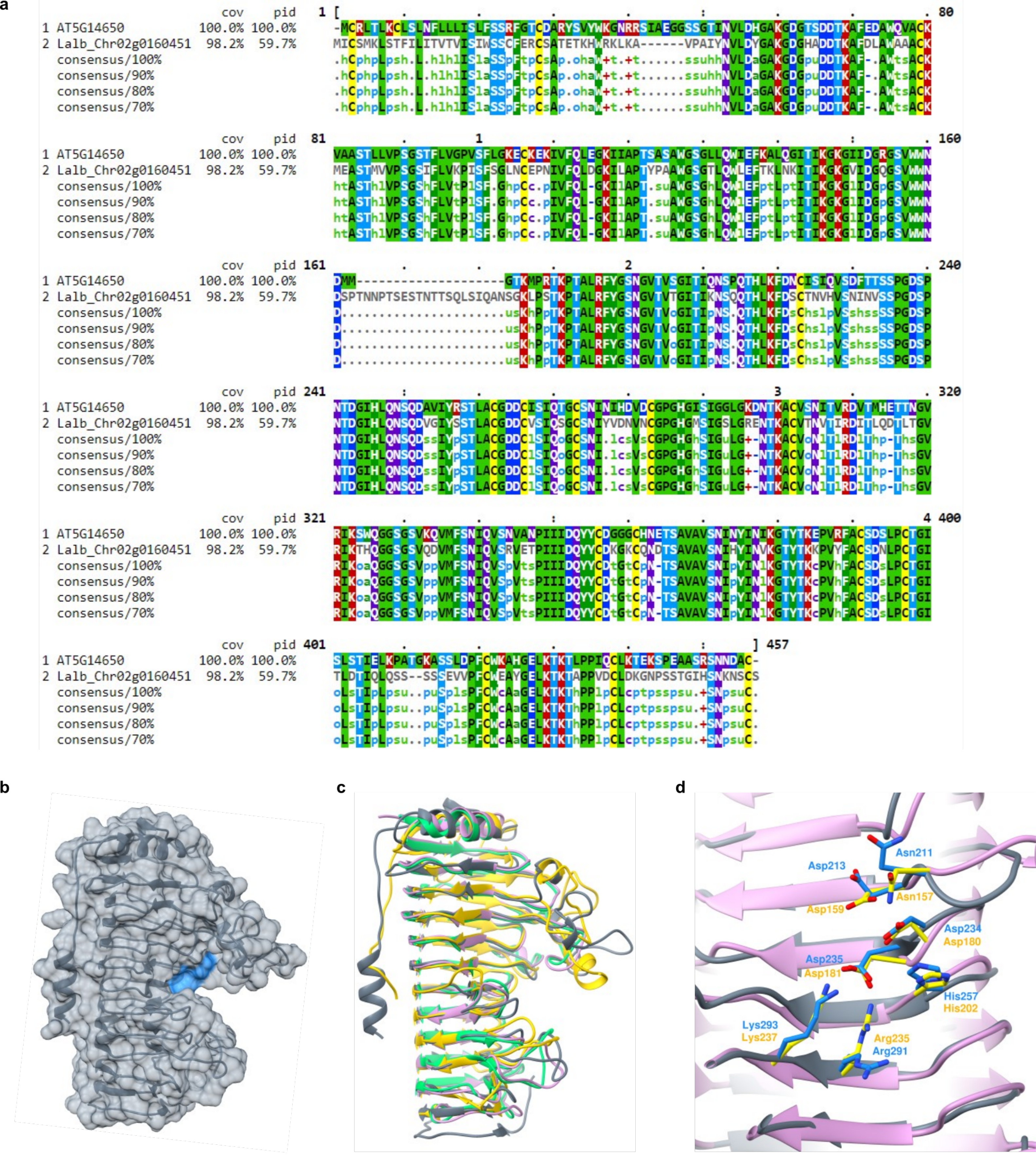


**Extended Data Fig.6: LaPOLYGALACTURONASE3 *in silico* characterization.**

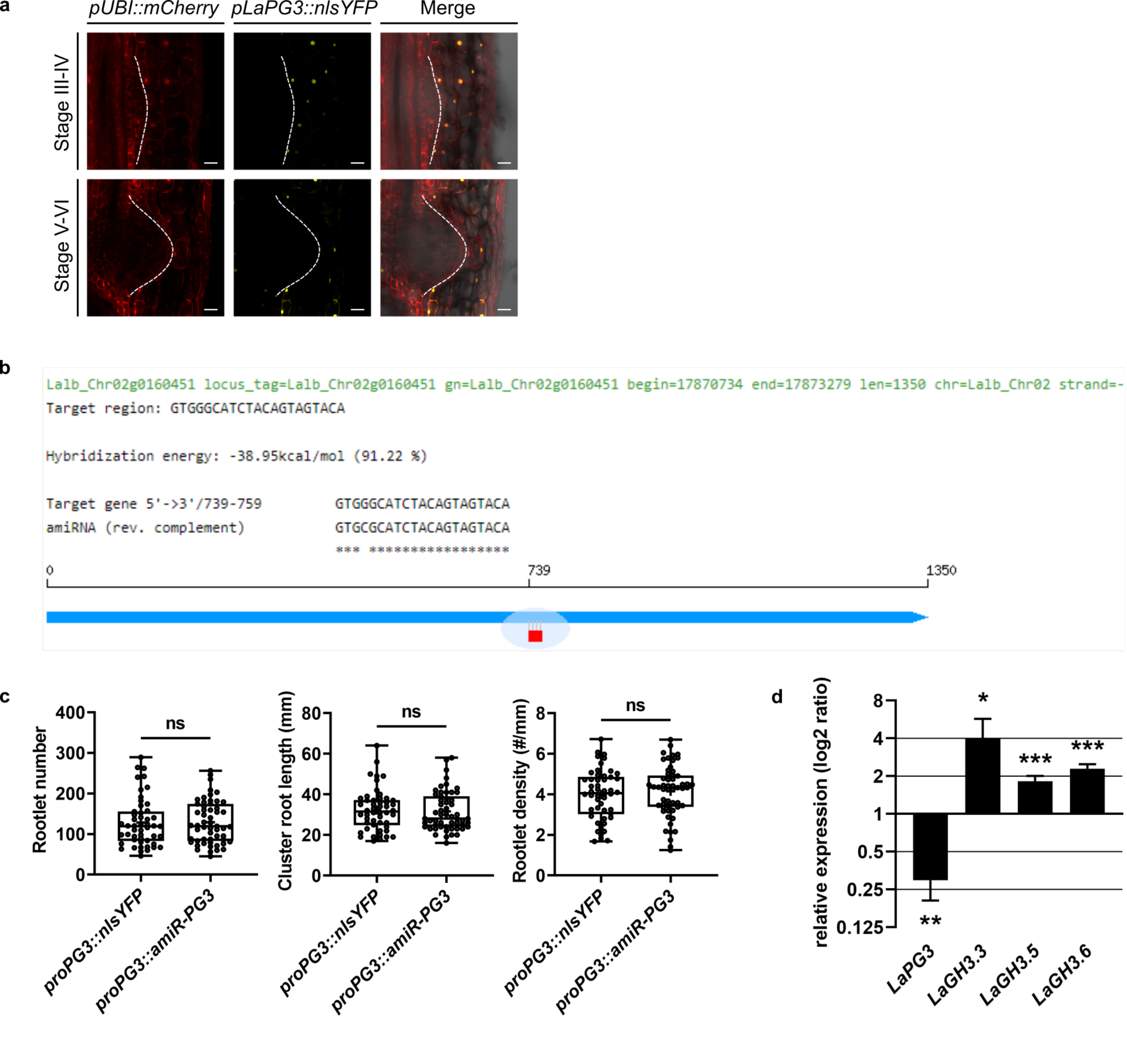


**Extended Data Fig.7: Characterization of *LaPOLYGALACTURONASE3* microRNA lines in white lupin hairy roots.**

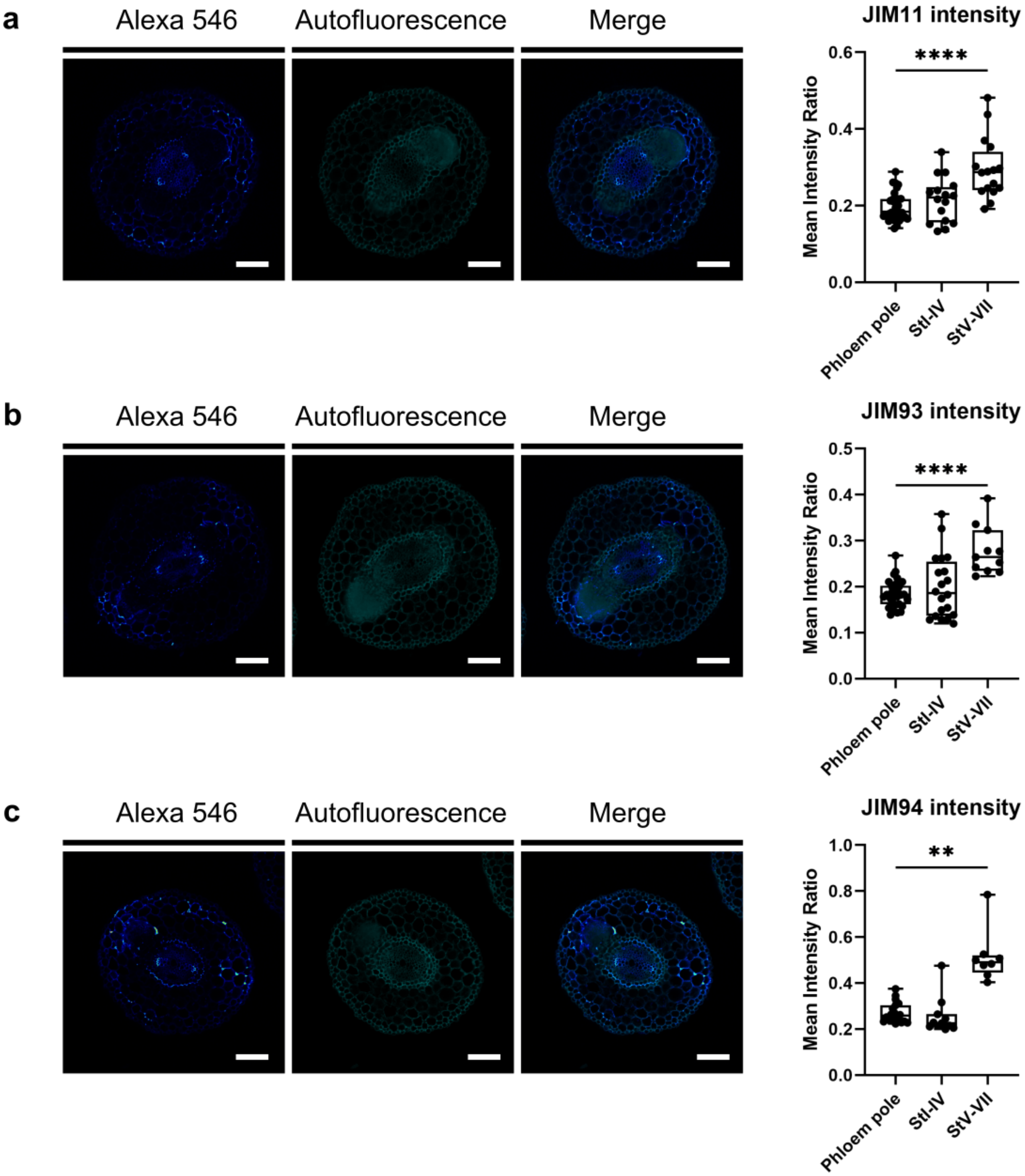


**Extended Data Fig.8: JIM11 and JIM93/JIM94 epitope labeling is increased in rootlet primordium-overlaying cells.**
